# Evolutionary rescue under demographic and environmental stochasticity

**DOI:** 10.1101/2023.03.13.532391

**Authors:** Kuangyi Xu, Todd J. Vision, Maria R. Servedio

**Affiliations:** Department of Biology, University of North Carolina, CB#3280, Coker Hall, University of North Carolina, Chapel Hill, NC, USA

## Abstract

Wild populations suffer two types of stochasticity: demographic stochasticity, from sampling error in offspring number, and environmental stochasticity, from temporal variation of the growth rate. By modeling evolution through phenotypic selection following an abrupt environmental change, we investigate how genetic and demographic dynamics, as well as effects of selection intensity after the environmental change and genetic variance on population survival, differ under demographic and environmental stochasticity. We find that the survival probability declines sharply with stronger selection under demographic stochasticity, but declines more continuously under environmental stochasticity. However, the genetic variance that confers the highest survival probability differs little under demographic and environmental stochasticity. Since the influence of demographic stochasticity is stronger when population size is smaller, a slow initial decline of genetic variance which allows quicker evolution and increase of fitness, is important for persistence. In contrast, the influence of environmental stochasticity is density-independent, so higher initial fitness becomes important for survival under strong environmental stochasticity. Combining both types of stochasticity shows that adding even weak environmental stochasticity can exaggerate the effects of different levels of demographic stochasticity on survival probabilities. Our work suggests the importance of explicitly distinguishing and measuring the forms of stochasticity for evolutionary rescue studies.

## Introduction

While genetic adaptation has long been a key focus of the study of evolution, it can obviously only occur given the prerequisite that the population does not go to extinction. In the past several decades, a substantial amount of research effort has therefore been put into the so-called framework of “evolutionary rescue”, which bridges both the genetic and demographic in the study of adaptation (see Martin 2013, Carlson et al. 2014, Alexander et al 2014, and Bell 2017 for reviews). Evolutionary rescue refers to the process whereby after an environmental stress, a population avoids extinction by genetic evolution (Gomulkiewicz and Shaw 2013), often with an initial demographic decline when the environmental change is abrupt, or is gradual but the initial rate of environmental change is too fast (Burger and Lynch 1995). Within this context, a great variety of models have been developed to investigate how genetic and ecological factors can influence population persistence (Carlson et al. 2014), and some of these theoretical predictions have been confirmed by studies in the lab (Martin et al. 2013).

Most of these theoretical studies focus on the probability that a population will survive versus go extinct, which use one of two major types of approaches. The first type focuses on the genetic changes that occur during adaptation. It usually assumes deterministic population growth and focuses on the stochasticity during genetic evolution. The population is considered rescued when adaptive alleles become fixed, and thus the survival probability can be calculated based on the fixation probability of the alleles (Orr and Unckless 2008, Orr and Unckless 2014, Matuszewski et al. 2015). The second type of model emphasizes demographic growth by allowing the population size to change with time. For simplicity, changes in the genetic composition of populations are approximated with deterministic equations. Some models of this type assume an extinction threshold below which extinction is likely to occur. the probability of extinction is generally assumed to increase with the time that the population stays under this threshold (e.g., Gomulkiewicz et al. 1995, Chevin and Lande 2010, Chevin et al. 2010, Osmond et al. 2013). These studies find that the survival probability deceases with the selection intensity and increases with the initial population size and genetic variance (Gomulkiewicz et al. 1995, Lynch et al. 1993).

Compared to the threshold population size assumption, more realistic models directly incorporate stochasticity during population growth, which allow expression of the population survival probability and mean time to extinction (e.g., Lynch and Lande 1993, Lande 1993, Gomulkiewicz et al. 2017, Anciaux et al. 2019). Two distinct types of stochasticity are often considered: demographic and environmental stochasticity. Demographic stochasticity is caused by sampling error during individual birth and death processes. In contrast, environmental stochasticity captures random fluctuations in the growth rate, which are often due to seasonal changes in reproduction, recruitment and survival probabilities. For example, in white-footed mice (*Peromyscus leucopus*) populations, recruitment is high during spring and low during autumn (Jacquot et al. 1998). In practice, population dynamics are often co-influenced by the two types of stochasticity (Stacey et al. 1992, Leirs et al. 1997, Grenfell 1998). Demographic stochasticity has been found to be more important in some populations (e.g., great tits, Sæther et al. 1998), while environmental stochasticity is more important in others (e.g., swiss ibex, Sæther et al. 2007; acron woodpeckers, Kendall 1998), depending on the specific species, life stages and habitats. Importantly, ecological models have shown that different types of stochasticity can lead to different predictions for the extinction probability. Generally, it is less likely for a single population to suffer extinction from demographic stochasticity than from environmental stochasticity (Lande 1993). However, in a prey-predator metapopulation, demographic noise is more likely to enhance extinction, while environmental noise may reduce interspecific interactions and thus contribute to coexistence (Engen et al. 1998).

Since demographic dynamics behave differently under the two types of stochasticity, key factors that determine whether a population will survive or go extinct in the evolutionary rescue process may also differ under the two types of stochasticity. Specifically, the effects on the survival probability of the selection intensity and genetic variance, two critical factors that affect evolutionary rescue in previous studies, should be assessed separately under demographic and environmental stochasticity. Previous meta-analyses show that the frequency of stabilizing selection intensity has a double exponential distribution, with most populations exhibiting weak selection during phenotypic evolution (Kingsolver et al. 2001). From an evolutionary rescue perspective, the paucity of records of strong selection may be due to a high extinction risk during the adaptation process. Stronger selection has opposing effects on rescue as it increases the rate of genetic evolution but also leads to more severe demographic decline (Haldane 1957, Alexander et al. 2014). Previous models show that the effects of selection intensity depend on the rate of environmental change. Under a sudden environmental shift, higher selection intensity tends to increase extinction risk (Gomulkiewicz and Holt 1995), while under a gradually moving phenotypic optimum, an intermediate selection intensity tend to be optimal for slowing down extinction (Burger and Lynch 1995). Genetic variance also has antagonistic effects during population adaptation. Genetic variance can be beneficial during evolutionary rescue because it enables rapid evolution. However, it also imposes a fitness cost under stabilizing phenotype selection (Lande and Shannon 1996). Indeed, previous studies show that effects of genetic variance depend on the severity and duration of environmental changes. When an environmental change is sufficiently long or extreme, larger genetic variance can promote population persistence, but a low level of genetic variance tends to be optimal for persistence if the change is temporary and short (Lyberger et al. 2021).

Here, by using stochastic differential equations, and adopting a deterministic approximation to the dynamics of the mean fitness, we compare how the population survival probability changes with selection intensity and genetic variance under demographic and environmental stochasticity. We focus on evolutionary rescue in a sexually reproducing population under stabilizing phenotypic selection following an abrupt environmental shift. To further determine the key factors that influence the fate of populations, we develop individual-based simulations incorporating both types of stochasticity and track the historical dynamics of genetic variance, mean fitness and population size across replicate populations. Recently, there has been growing interest in tracking the survival probability of populations over time, since most empirical observations cover a short time span (Martin et al. 2013, Gomulkiewicz et al. 2017, Anciaux et al. 2019). A detailed study of the distribution of extinction times after an environmental shift shows that populations with a higher survival probability than others in early generations may suffer a higher extinction risk later on (Bell 2017). Therefore, throughout the paper, we focus on the temporal dynamics of the rescue process. We find that the effects of selection intensity on survival probability differ under demographic and environmental stochasticity, while the optimal level of genetic variance that gives the highest survival probability is robust to types of stochasticity. Under demographic stochasticity, it is crucial to have a slower decline of genetic variance for a population to survive later generations of the rescue process, while under environmental stochasticity, a high average fitness in the initial generations after an environmental shift is what is predictive of successful evolutionary rescue. For populations under both types of stochasticity, adding even weak environmental stochasticity can exaggerate the effects of different levels of demographic stochasticity on the survival probability.

## Methods

### Model framework

We consider a sexually reproducing, random mating population under stabilizing phenotypic selection that suffers an abrupt environmental change. Individual fitness depends on the phenotypic value of a quantitative trait, and there is an optimal phenotypic value that gives the highest fitness at a certain environment. Initially, the population consists of *N*_0_ individuals assumed to reach the selection-drift-mutation balance. After a sudden environmental change, the optimal phenotypic value shifts, and selection then causes the population to evolve towards the new optimum. The population suffers a risk of extinction due to demographic and environmental stochasticity during the evolutionary rescue process.

### Selection and fitness

We assume the population is subject to stabilizing selection on a quantitative trait. The fitness landscape is modeled as a quadratic function with a single phenotypic optimum (Lande and Shannon 1996, Anciaux 2019). Initially, the phenotypic optimum is at the value *z*_0_, which gives the highest fitness. After the environment changes, the optimum shifts to a new value at *z*_0_ + *d*_0_, where *d*_0_ measures the level of the shift. Without loss of generality, we can set *z*_0_ = 0. In the new environment, the fitness of individuals with phenotype *z_i_* (in the sense of Malthusian parameter) is given by:

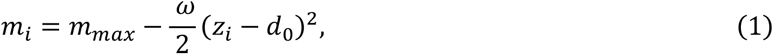

where *m_max_* is the maximum fitness at the phenotypic optimum, and *ω* measures the selection intensity.

### Stochastic demographic growth

To approximate continuous population growth, we adopt the method of stochastic differential equations (SDE), which is used extensively in previous work (e.g., Turelli 1977, Lande et al. 1993, Gomulkiewicz et al. 2017, Anciaux et al. 2019). We assume the population growth is density-independent, and the noise is not autocorrelated. By averaging the fitness across all individuals in the population, we show in Appendix I that the dynamics of the total population size *N_t_* can be written as:

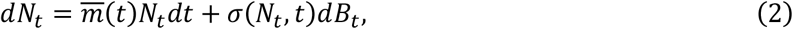

The first term describes deterministic growth, where 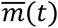 is the average per capita growth rate across the population. The second term accounts for stochasticity, where *B_t_* is the Wiener process, and *σ*^2^(*N_t_, t*) is the variance of stochasticity, which can depend on both time and the population size.

We consider two types of stochasticity: demographic and environmental stochasticity (Ovaskainen et al. 2010). For demographic stochasticity, 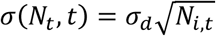, and for environmental stochasticity, *σ*(*N_t_, t*) = *σ_e_N_t_*, where 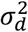 and 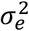 are the strength of demographic and environmental stochasticity, respectively. Note that the environmental stochasticity modeled here is different from that in Chevin et al. (2017), where they define environmental stochasticity as random fluctuations in the phenotypic optimum. During a time period *dt*, the variance of the relative population size change *Var*(*dN_t_/N_t_*) is 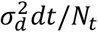 and 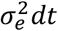 under demographic and environmental stochasticity, respectively. Therefore, the influence of demographic stochasticity gets stronger as the population size becomes smaller, while the influence of environmental stochasticity is independent of the population size. The condition for extinction differs between the two types of stochasticity. Under demographic stochasticity, a population is considered to go extinct when *N_t_* = 0. However, for environmental stochasticity, we assume that extinction happens when the population size decreases to an extinction threshold *N_t_* = 1, since with only environmental stochasticity, the population size will never go to 0 in the SDE model. As for the values of 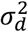 and 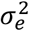, previous studies have shown that the demographic stochastic strength 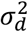 is usually at the order of ~ 1, while 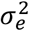 is often estimated to be smaller than 0.1 (Kendall 1998, Lande et al. 2006, Sæther et al. 2007). However, in Appendix II, we show that for populations under evolutionary rescue which suffer demographic decline, a larger value of 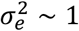 is possible.

### Fitness dynamics

We assume a Gaussian distribution for the phenotype frequency with a mean value 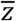 and a phenotypic variance 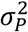, which is the sum of genetic variance 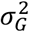 and the environmental variance 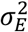. Integrating the phenotypic frequency distribution by its fitness, the population growth rate is 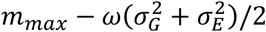, where the second term shows the fitness reduction caused by genetic and environmental variances (Lande and Shannon 1996). During the evolutionary rescue process, we assume that both the genetic and environmental variance are nearly constant, although the genetic variance may be reduced under strong selection (this is varied in the section of individual-based simulations). Based on the assumption, the population growth rate 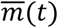 after an environmental shift *d*_0_ is (Equations 2 and 4 in Lande and Shannon 1996)

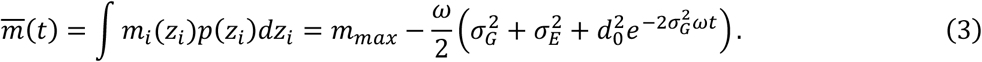

The third term within the parentheses represents the fitness cost imposed by the environmental shift, which decreases exponentially with time due to genetic evolution. Note that this equation ignores stochasticity in mutation, drift and recombination, which may not hold when the population size becomes small.

### Survival probability

When a population is only subject to demographic stochasticity, the demographic dynamics follow the SDE:

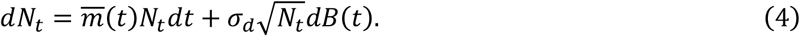

Given that a population starts with an initial size *N*_0_, the probability that a population is still viable before time *t* is (Anciaux et al. 2019):

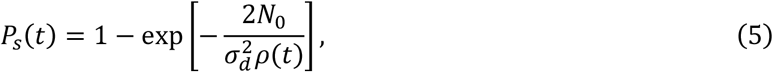

where

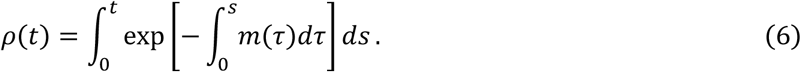

This expression helps us separate the effects of several key component factors on the survival probability. The population survival probability increases with its initial size *N*_0_, while it decreases with the strength of stochasticity 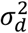. The integral *ρ*(*t*) captures the evolutionary trajectory of the population mean fitness, which is determined by both the extrinsic selection intensity and intrinsic genetic factors. From direct inspection of the expression of *ρ*(*t*), the survival probability is mostly determined by fitness dynamics in the initial generations, as the contribution of 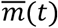 to the value of the integral of *ρ*(*t*) decreases exponentially with time.

### Individual-based simulation

In the above model, genetic variance is assumed to be constant during evolution, and we also ignore other factors like genetic drift. In order to relax these assumptions, we also performed an individual-based simulation written in C++ (the code is available on Dryad [submitted upon acceptance]). Following previous work (Boulding et al. 2001, Anciaux et al. 2019), we assume non-overlapping generations. Survival probability is estimated by running 500 replicate simulations starting with the same population. A simulation stops when a population goes extinct, is considered to be rescued, or reaches the maximum time (usually 100 generations since we focus on a short time scale). A more detailed description and discussion of the simulation algorithms can be found in Appendix IV.

Briefly, our simulation mainly consists of two parts: the genetic and demographic parts. For the genetic part, we adopt the method from Boulding et al. (2001) to simulate the genetic basis of phenotype. We assume a random mating population with diploid, hermaphroditic individuals. Each individual has a pair of two chromosomes, and their phenotypes are controlled by *N_s_* identical biallelic loci located on the chromosome that act additively with equal forward and backward mutation rates. To eliminate the effects of the number of loci *N_s_*, the phenotype effect of each allele is scaled according to *N_s_*, and. We assume free recombination between loci. To maintain the genetic variance, we assume a constant genomic mutation rate *U* every generation, so that the forward and backward mutation rates at each locus are *U*/2*N_S_*. Although we assume hermaphroditic individuals, the results should also apply to populations with separate sexes. The major difference between the two types of populations is that hermaphrodites can self-fertilize. However, we assume random mating, so that selfing is rare except in very small populations, where stochasticity will be a much larger factor. The survival probability is expected to be similar under hermaphroditism and the case of separate sexes provided that the number of females is equal to the number of hermaphrodites.

The algorithm for the demographic portion of the simulation differs between demographic and environmental stochasticity. Individual fitness *m_i_* can be calculated based on their phenotype using equation (1), from which we can obtain the average growth rate 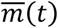 and the mean absolute fitness 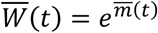. The population size in the next generation is drawn from a Poisson distribution 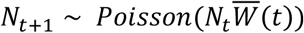, which makes the strength of demographic stochasticity 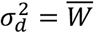. For the simulation of environmental stochasticity, in each generation, we add a random number *δ* drawn from the normal distribution 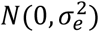 to the mean growth rate 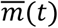, and the absolute fitness becomes 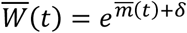. Since the offspring number 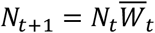 is a numeric value, to make it an integer, we adopt two functions: 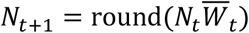, and 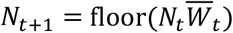, but the two functions yield very different results for the survival probability (Fig. S1). This is because when the population size shrinks to an extremely low level, even a difference of only one individual can greatly influence the ultimate fate of a population (Fig. S2, and see Appendix IV for a detailed explanation). This intrinsic limitation of the simulation makes it difficult to compare results from the analytical model with simulations when populations are under environmental stochasticity.

## Results

### Effects of the selection intensity on the population survival probability differ under demographic and environmental stochasticity

The selection intensity has antagonistic effects on population survival. In equation (3), stronger selection results in a higher fitness load under stabilizing selection, but also leads to quicker evolution. Since different traits and populations have different phenotypic variances, to allow comparisons, the selection intensity is measured in terms of the fitness decrease in units of phenotypic variance 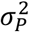. Without loss of generality, we can set 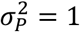, and the fitness in equation (3) can be written as:

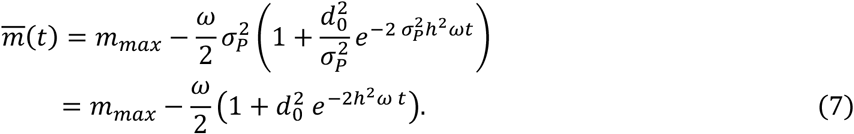

where *d*_0_ is also measured relative to the phenotypic variance 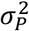, and 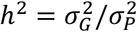 is the heritability. From equation (7), we can calculate two critical values of selection intensity, *ω*_1_ and *ω*_2_, which respectively make 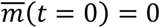 and 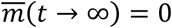. When *ω* < *ω*_1_, the initial fitness is positive after the environmental shift, so the expected growth rate of a population is always positive during the adaptation process. When *ω* > *ω*_2_, the average growth rate of the population is expected to be always negative throughout the adaptation process.

When the selection intensity is intermediate (*ω*_1_ < *ω* < *ω*_2_), the population will have a negative initial growth rate 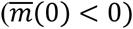 and a positive one after genetic evolution 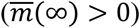. This means that the population size is expected to first decrease and later increase, so that the demographic change is characterized by a U-shaped curve, which is the classical situation considered in previous studies of the evolutionary rescue process (Gomulkiewicz and Holt 1995, Gomulkiewicz1 et al. 2012).

When the population is subject to demographic stochasticity, we can qualitatively see how selection intensity influences the survival probability, based on equation (5). It can be proven that at any time *t*, stronger selection always lowers survival probability, as:

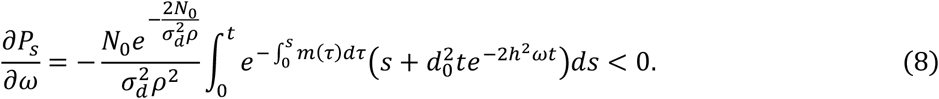

However, the effects of selection intensity on the survival probability depend on time and differ between the two types of stochasticity. Fig. 1 illustrates the changes of survival probability with selection intensity at different time points under demographic and environmental stochasticity. Depending on the selection intensity *ω*, the three sections correspond to the three situations described in the last paragraph: the growth rate is always positive (section I), there is a U-shaped demographic curve (section II), and the growth rate is always negative (section III).

**Figure 1.**
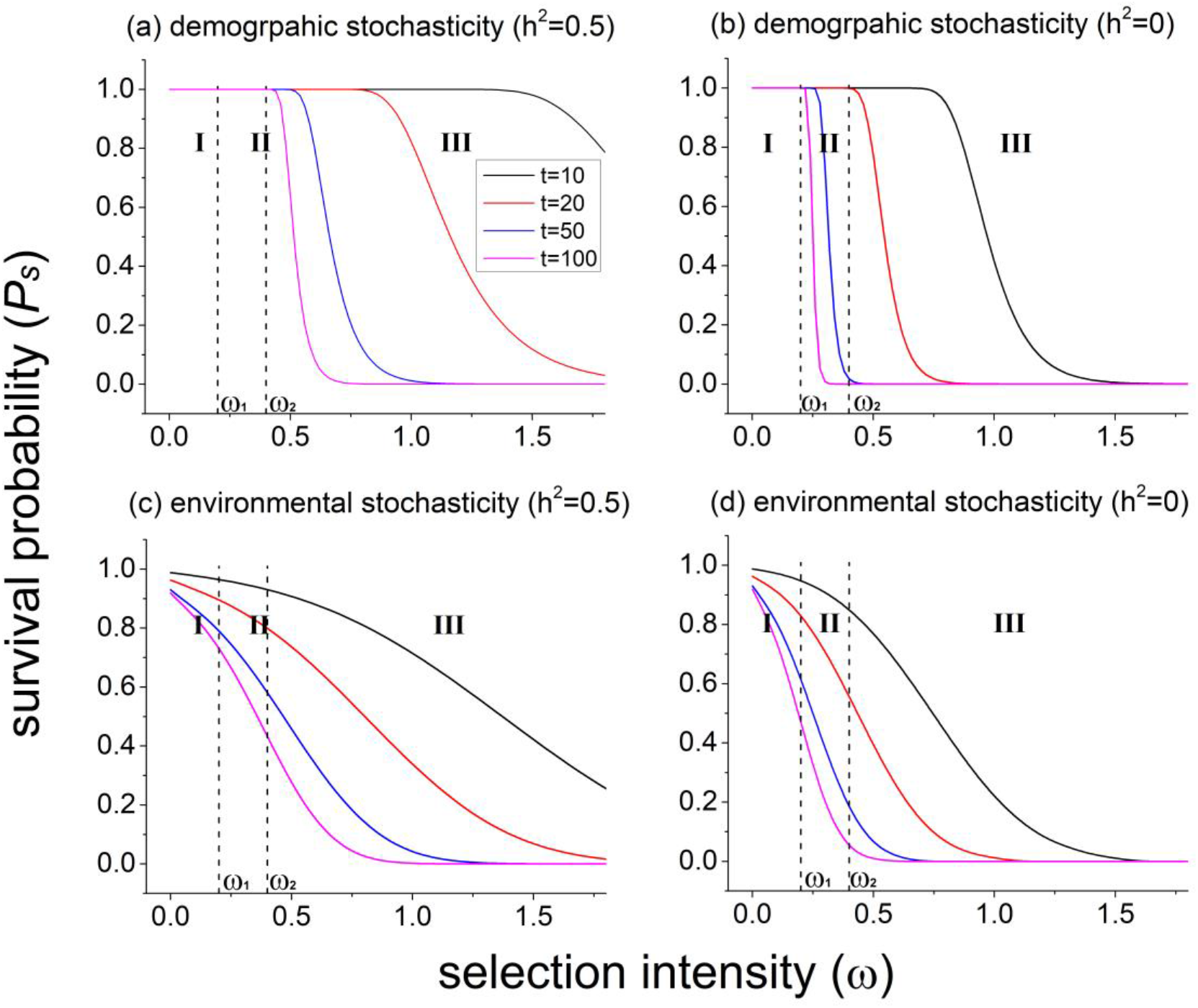
Changes in the survival probability over the selection intensity at certain time points (*t* = 10,20,50,100) under demographic and environmental stochasticity with genetic evolution *h*^2^ = 0.5 and no genetic evolution (*h*^2^ = 0). The values of the selection intensity at the two dashed lines are *ω*_1_ = 0.2 and *ω*_2_ = 0.4 from left to right respectively. Populations with a strength of selection smaller than *ω*_1_ (section I) will always have 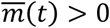 during the whole evolutionary rescue process, while the growth rate is always negative 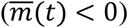 for those in section III (*ω* > *ω*_2_). In the interval between *ω*_1_ and *ω*_2_ (section II), the population growth rate changes from negative to positive, which is the typical case of evolutionary rescue considered in previous work (the U-shaped curve, Gomulkiewicz et al. 1995). Parameter values are *N*_0_ = 500, *d*_0_ = 1, *m*_0_ = 0.2, 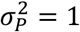, with 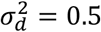 for panels (a), (b), and 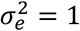 for panels (c), (d).

Under demographic stochasticity, as shown in Figs. 1(a) and 1(b), at the beginning of the rescue process (*t* = 10), extinction events only happen in populations subject to strong selection (*ω* > 1.3 in Fig. 1(a) and *w* > 0.8 in Fig. 1(b)). After a time period of about 50 generations, the decrease of the survival probability becomes quite sharp, with most changes of *P_s_* only lying within a narrow range of *ω*, and the range shifting towards the left over time. Moreover, when there is substantial heritability and thus quick phenotypic evolution (Fig. 1(a)), the decline of the survival probability mainly happens in section III, in which populations always have a negative growth rate, while in sections I and II, populations retain a survival probability of nearly 100%. In contrast, without phenotypic evolution (*h*^2^ = 0, Fig. 1(b)), differential survival probabilities mainly occur in section II. Therefore, the typical scheme where the population size initially decreases and then subsequently increases (the U-shaped demographic curve) may not always be a suitable framework for the study of evolutionary rescue in the context of an abrupt environmental change.

In contrast, under environmental stochasticity, we do not see a steep decline of the survival probability with the strength of selection, as we do under demographic stochasticity. As shown in Figs. 1(c) and 1(d), throughout the period of 100 generations, the decrease of population survival probabilities spans across the three sections I, II and III. Unlike that under demographic stochasticity, even at the early stages of evolutionary rescue (*t* = 20), a decrease in the survival probability can be found at small *ω*. Moreover, the range of *ω* for which the survival probability initially decreases does not shift with time. These patterns are an intrinsic characteristic of environmental stochasticity, as even under weak selection, populations with a positive growth rate are still at risk of extinction due to fluctuations in mean fitness. Once again, under environmental stochasticity, the extinction events span across all three sections. Instead, our results suggest that evolutionary rescue is a relevant process even when there is initially a positive growth rate after an environmental shift; the population still suffers an elevated extinction rate in this case (region I), which can be reduced by evolution towards the new optimum. These results therefore call for a broadening of the perspective of when the concept of evolutionary rescue is relevant (away from a requirement of a U-shaped curve).

### Effects of genetic variance on survival probability

Here we investigate the effects of genetic variance under two scenarios: populations have the same phenotypic variance 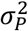, or the same environmental variance 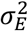. Biologically, when comparing populations distributed across different habitats, it is more suitable to discuss the effects of genetic variance under the first scenario. On the other hand, we will expect to see the second situation for populations living in the same or similar habitats, where they experience almost the same developmental stochasticity and environmental fluctuations.

#### Populations have the same phenotypic variance

When the phenotypic variation 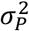 is fixed, a higher proportion of genetic variance (i.e., heritability) will lower the extinction risk, since populations benefit from a faster rate of evolution towards the new optimum. However, it is of empirical interest to know to what extent that a higher genetic variance can enhance the population survival probability. To empirically detect the effects of genetic variance on the survival probability, a significant increase of survival probability spanning a wide range of genetic variance would be needed. Here we show that the level of this increase is contingent on the strength of selection and on time. Under the assumption that the phenotypic variance is fixed, we set *σ_P_* = 1 and use heritability *h*^2^ as an indication of the level of genetic variance 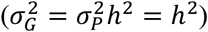, as in the previous section.

Fig. 2 shows changes of the survival probability over the spectrum of heritability *h*^2^ under demographic stochasticity. Generally, the change of the survival probability behaves in a logistic manner as heritability increases. As indicated in Fig. 2(a), an intermediate level of selection intensity (*ω* = 0.5,0.8) is more likely to reveal the effect of the genetic variance, resulting in a survival probability curve spanning across the whole spectrum of heritability. However, under the same selection intensity, the change of survival probability with heritability also depends on time. For *ω* = 0.8, the survival probability changes the most with heritability at the beginning of the rescue process (*t* = 10), with only little dependency in later generation (i.e., at *t* = 50). In contrast, under weak selection (*ω* = 0.1), only over the longer term (*t* = 50) do we find that the genetic variance has the most prominent effect on enhancing the survival probability. Practically, to detect the effects of genetic variance on survival probability, the time at which the observations are made should be carefully chosen. In order to reliably detect differential survival probability between populations with different levels of genetic diversity, an early observation is preferred if selection is strong, while it would be better to wait longer if selection is weak. This conclusion also holds for populations under environmental stochasticity (see Fig. S3).

**Figure 2.**
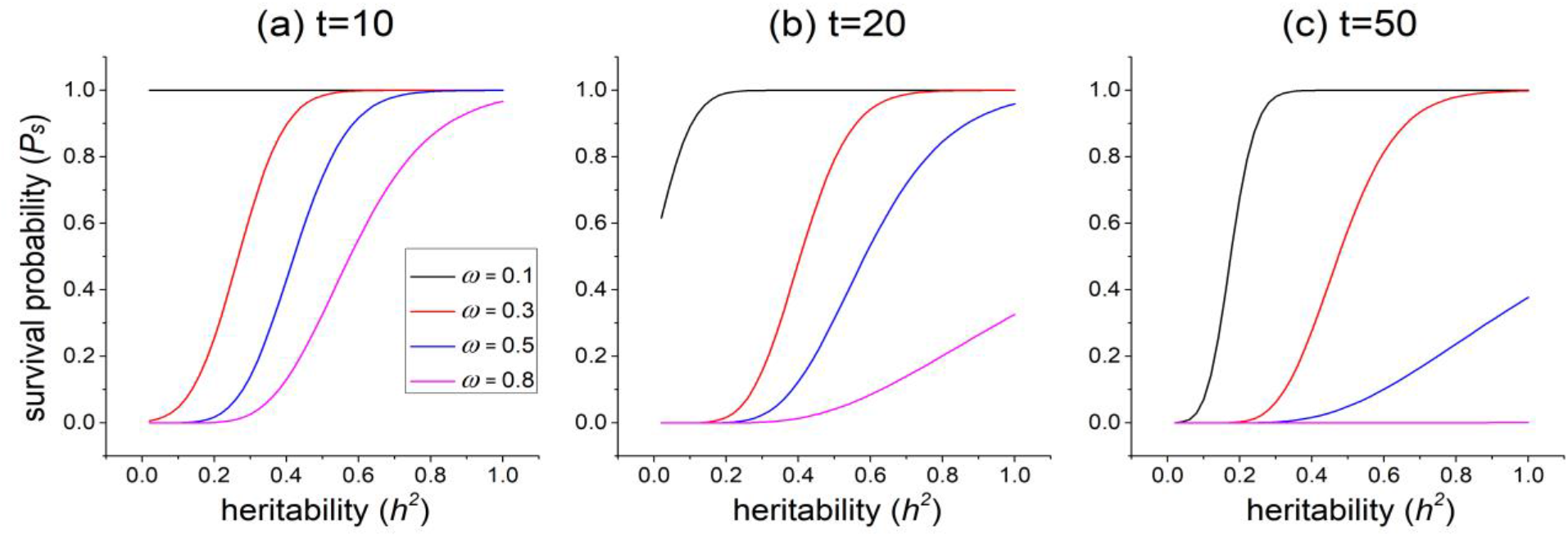
Changes of the survival probability with heritability *h*^2^ under demographic stochasticity, given fixed phenotypic variance. Panels (a), (b), and (c) show snapshots at the three time points *t* = 10,20,50 espectively. In each panel, the colored lines indicate a different selection intensity at *ω* = 0.1,0.3,0.5, and 0.8. Generally, the most prominent change of over *h*^2^ happens earlier in time and under stronger selection. For *ω* = 0.8, the most prominent change appears at time *t* = 10, while for *ω* =0.1, heritability has the most significant effects on population survival at *t* = 50. Parameters for all three panels are 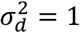, *N*_0_ = 500, *d*_0_ = 3, *m*_0_ = 0.2, and 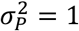.

#### Populations have the same environmental variance

When the environmental variance 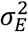 is fixed, we can set 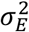 to be the unit of measurement so that 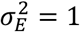. The mean growth rate in equation (3) becomes:

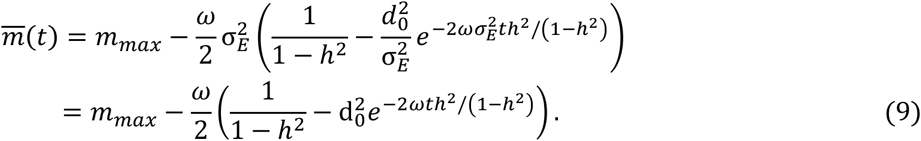

Here 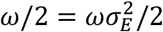 directly measure the fitness load imposed by the environmental variance, and the shift of the phenotypic optimum *d*_0_ = *d*_0_/*σ_E_* is now measured relative to *σ_E_*. We again choose the heritability *h*^2^ as a measurement of the level of genetic variance. However, in this case 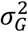 is no equal to *h*^2^ but rather 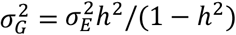.

Given a fixed level of environmental variance, a larger genetic variance does not always ensure a higher survival probability. This is because genetic variance can impose a fitness load under stabilizing selection, although it increases the evolution rate (Lande and Shannon 1996). During the evolutionary rescue process, these two antagonistic effects compete with each. Fig. 3 illustrates how heritability influences the dynamics of the population survival probability under demographic stochasticity. At each time point, there exists a certain level of heritability that gives the highest survival probability, which we call the “heritability optimum”, which is not invariant but depends on the amount of the environmental shift *d*_0_ and also changes with time *t*. When the environmental shift is not large, populations do not have to evolve quickly to catch up with the new optimum. Therefore, populations with low levels of genetic variance are favored because they have a small fitness load. In Fig. 3(a), when there is only slight change in the phenotypic optimum (*d*_0_ = 0.5), populations with no genetic variance 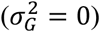 have an advantage in the early stage before 20 generations, and only later does the heritability optimum slowly increase to a higher level over time. However, it should be noted that when the environmental shift *d*_0_ is small, even populations with a genetic variance far from the optimum can still maintain a high survival probability, as the survival probability is higher than 75% in Fig. 3(a) as long as the heritability is below 0.5. As the environmental shift gets larger, the heritability optimum quickly increases. When *d*_0_ = 1, populations with *h*^2^ = 0 only have an advantage over others for a few generations. As the level of the shift becomes quite large (*d*_0_ = 3), the heritability optimum starts at a high level and gradually decreases with time.

**Figure 3.**
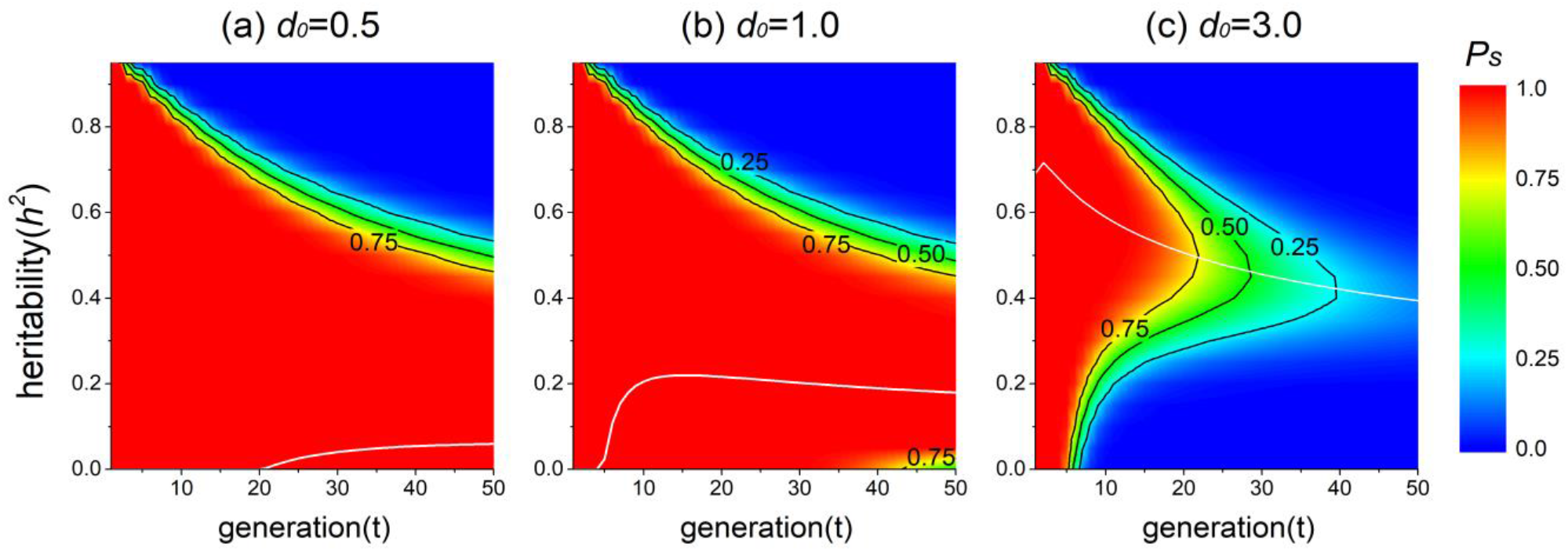
The survival probability landscape across different levels of heritability under demographic stochasticity given a fixed environmental variance. Results are calculated from equation (5). The three panels (a), (b) and (c) show the survival probability under the level of environmental shift at *d*_0_/*σ_E_* = 0.5,1, and 3 respectively. The white solid line in each panel marks the heritability optimum. Under small *d*_0_ = 0.5, the heritability optimum stays at *h*^2^ = 0 until about *t* = 20, when it begins to increase. Under a larger environmental shift (*d*_0_ = 3), however, the heritability optimum quickly rises to a high value of about *h*^2^ = 0.7 and then decreases and converges to a lower level over time. Other parameter values are 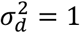, *N*_0_ = 500, *ω* = 0.3, and *m*_0_ = 0.2.

The shape that the survival probability changes with heritability and time under environmental stochasticity differs from that under demographic stochasticity (Fig. 4), and it may be expected that the heritability optimum may be influenced by different types of stochasticity. Nevertheless, we find that the value of the heritability optimum is nearly the same under environmental stochasticity or a combination of both types of stochasticity, as can be seen by comparison of the white lines in Fig. 4 and Fig. 3(c).

**Figure 4.**
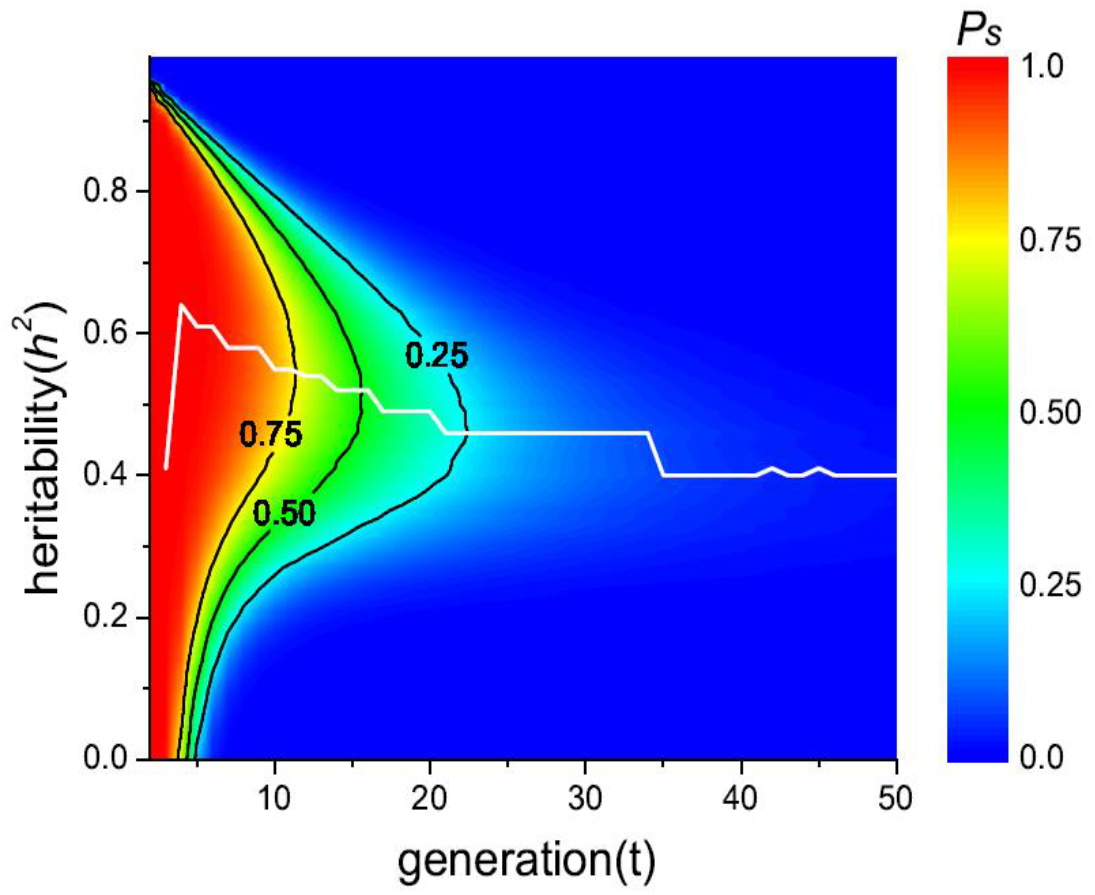
The survival probability landscape across heritability under environmental stochasticity given a fixed environmental variance 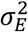. The white line indicates the heritability optimum. Results are computed by numeric solution of equation (A4.1), with the mean growth rate modeled by equation (8). Parameters used are 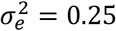, *N*_0_ = 500, *d*_0_ = 3, *m*_0_ = 0.2.

### Key factors that determine the fate of a population under demographic and environmental stochasticity

During the evolutionary rescue process, both genetic and demographic properties of the population vary with time due to factors like drift, selection and mutation. We expect that the trajectories of the genetic and demographic properties of a population at the initial stage of the rescue process may bear some relationship to survival or extinction in the later stage. We use simulations to investigate the key factors that determine the ultimate fate of populations when considering both genetic and demographic dynamics together.

Populations under demographic stochasticity can successfully survive the rescue process because they fortunately have the chance to maintain a relatively higher level of genetic variance than those that go extinct. We divide replicate populations from our individual-based simulations into two groups based on whether they go extinct or survive within 100 generations. Although there are some stochastic fluctuations, genetic variance generally drops in both groups after the environmental shift (Fig. 5(a)), mainly due to the action of strong selection and an increased level of drift after the quick decay of population size. However, those populations that avoid extinction tend to have a relatively minor decline of genetic variance in the initial 20 generations (red line in Fig. 5(a)). This difference in the dynamics of genetic variance is quite important because it benefits the survival group by allowing more rapid evolution. As shown in Fig. 5(d), the mean growth rate of the survival group evolves more rapidly to a positive value than in the extinction group.

**Figure 5.**
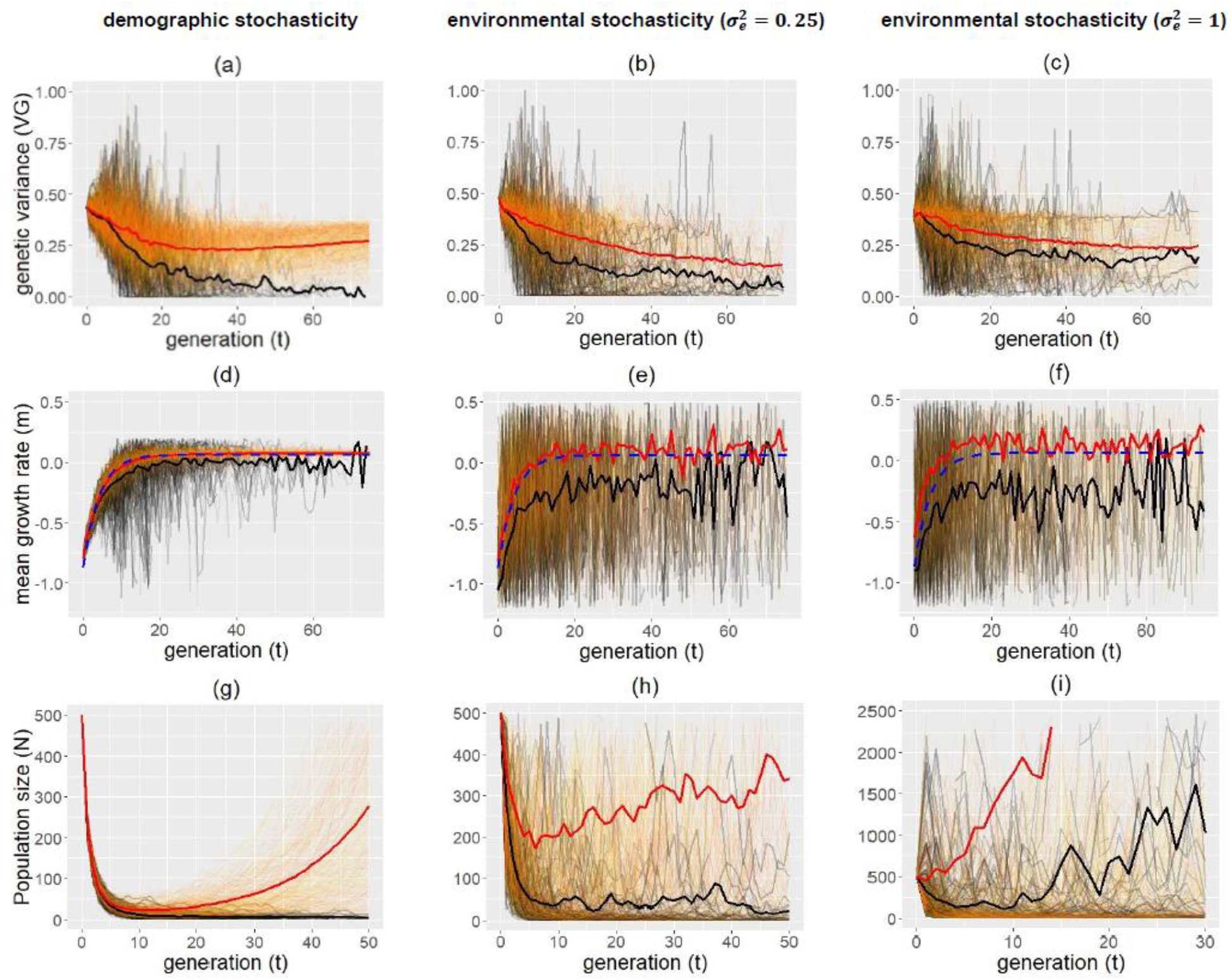
Changes of genetic variance, the mean growth rate and population size with time across 500 replicate populations in three cases. In all panels, populations that go extinct within 100 generations (extinction group: 205, 195 and 259 replicates populations under demographic stochasticity and environmental stochasticity with 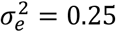 and 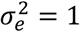, respectively) are colored from grey to black, while those that survive through the time period (survival group) are colored from yellow to red. The average genetic variance, growth rate and population size across the extinction and survival groups are plotted by the black and red bold lines respectively. In panels (d)-(f), the blue dashed line depicts the fitness predicted from equation (3) by plugging the initial value of genetic variance into the model. For demographic stochasticity, the offspring number is drawn from the Poisson distribution with mean 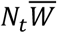, so that 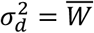. Simulation of environmental stochasticity used the round function. Other parameter values used in the simulation are that *N*_0_ = 500, *m_max_* = 0.2, *ω* = 0.3, *d*_0_ = 2.5, *V_P_* = 1, *V_E_* = 0.5, and *U* = 0.5.

Although the fate of populations is related to the temporal dynamics of genetic variance, our SDE model, which only captures stochasticity in the demographic aspects, seems to predict the survival probability well, because there is a correlation between the population size and the change of genetic variance. In Fig. 5(g), although the size of both groups quickly drops to a low level within a few generations after the environment shifts, the survival group on average has a slightly higher population size than the extinction group. However, as stochasticity happens in both the demography and the genetic variance, the direction of causality is unclear. In other words, 1) does the larger population size result from higher fitness caused by a higher genetic variance; or 2) does the higher genetic variance result from lower genetic drift due to a large population size; or 3) do these processed mutually interact with one another? Additionally, for the survival group, there is an increase of genetic variance in later generations (after about 30 generations) in Fig. 5(a), which is due to the accumulation of new mutations after the population size becomes large. In contrast, the extinction group quickly loses their genetic diversity over a few generations, which results in a slow evolution rate at the beginning of the process. Moreover, the prospect for a population to survive declines even further in later generations, because strong drift due to the decline in population size can quickly fix alleles and further decrease genetic diversity, resulting in an even slower evolution rate.

Under strong enough environmental stochasticity 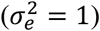, successful adaptation occurs when populations have the good luck to happen to have a higher growth rate. As illustrated in Fig. 5(c), there is not much difference in the genetic variance between the extinction and survival groups. Although the survival group does have a slightly higher level of genetic variance in the initial several generations, this is not a major cause of its survival but is more a result of a larger population size due to higher fitness. The two groups have a similar rate of fitness increase (the red and black lines are parallel in Fig. 5(f)), which indicates that the slight difference of genetic variance plays a minor role. Maintenance of a low level of genetic variance is due to a small effective population size resulting from frequent fluctuations in the population size. As Fig. 5(f) shows, despite fluctuations, the average fitness across the survival group is much higher than that of the extinction group in early generations. The advantage of this higher fitness can be seen from the dynamics of population size. In Fig. 5(i), unlike in the case of demographic stochasticity, there is only a slight initial decline in the average size of the survival group under environmental stochasticity. Under environmental stochasticity, genetic variance no longer plays a key role in determining the fate of populations.

The above differences under the two types of stochasticity can be intuitively understood by noting that the variance of the relative population size change is 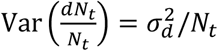 under demographic stochasticity and 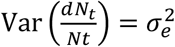 for environmental stochasticity. Demographic stochasticity causes positive feedback as population size gets smaller, and a quicker initial evolution (thus a slower population size decline) allows a population to escape from extinction (Nabutanyi and Wittmann 2021). In contrast, environmental stochasticity can still play a role even when population size grows to be large, and when the strength is strong enough, it can obscure the effects of rate of adaptation. When strength of environmental stochasticity is moderate 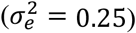, the genetic and population size dynamics behave at the intermediate level between that under purely demographic stochasticity and strong environmental stochasticity (see panels with 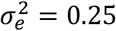 in Fig. 5), and both genetic variance and a higher growth rate due to stochasticity can play a role (see Figs. 5(b) and 5(e)).

It is expected that the strength of environmental stochasticity may depend on the absolute fitness of the population 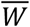. Therefore, we further investigate a special case when the strength of environmental stochasticity is proportional to the absolute fitness (i.e., 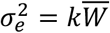, where *k* measures the strength of dependency). In this case, 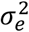 will be small at the initial stage, and gradually increases as fitness recovers. Generally, the pattern is similar to that under a constant strength of environmental stochasticity (Fig. S4), but the population size of the survival group size cannot recover quickly.

### Joint effects of demographic and environmental stochasticity on survival probability

In previous sections, the effects of the two different types of stochasticity on survival probabilities are analyzed separately. Here we combine both types of stochasticity to see how they co-influence the population survival probability. In Fig. 6, the survival probability drops as the level of environmental stochasticity increases (Boulding et al. 2001). The two types of stochasticity do not act in an additive way on the survival probability, that is, decrease of the survival probability under both types of stochasticity is often smaller than the sum of the decreases when only one type of stochasticity is present. Interestingly, the survival probability under both types of stochasticity together is also lower than the product of the survival probabilities under each type of stochasticity individually, suggesting a less-than-multiplicative interaction.

**Figure 6.**
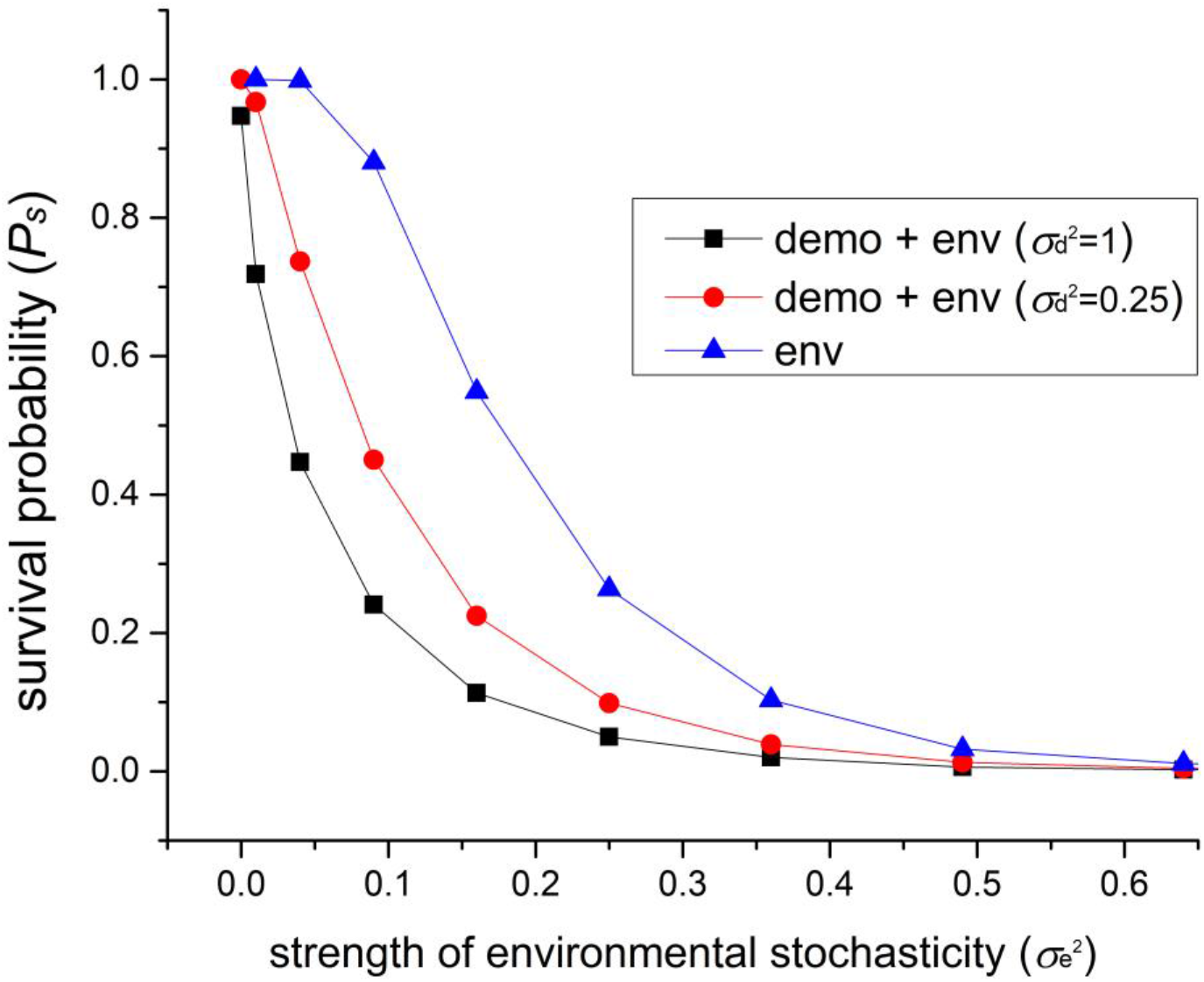
The population survival probability under both demographic and environmental stochasticity. The figure shows how the survival probability at *t* = 50 decreases with the strength of environmental stochasticity 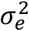. The results are computed by numeric solution of the SDE equation (A3.3) (see Appendix III). For each line, the strength of demographic stochasticity is fixed, with 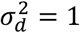 for the black line, 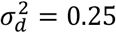 for the red line, and 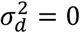 for the blue line. Therefore, the blue line shows populations with only environmental stochasticity, and the two data points at 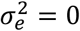 represents populations with only demographic stochasticity. The survival probability decreases with the strength of environmental stochasticity in a non-additive manner. Other parameter used in the results are that *N*_0_ = 500, *N_e_* = 1, *m_max_* = 0.2, *ω* = 0.3, *d*_0_ = 2.5, 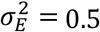, *U* = 0.5.

More importantly, the existence of a certain level of environmental stochasticity can reveal the effects of different levels of demographic stochasticity on the population survival probability. When there is only demographic stochasticity present, different levels of 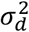 give similar survival probabilities (compare the two points at 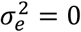 in Fig. 6). However, the difference in survival probability under different levels of demographic stochasticity increases after adding some environmental stochasticity. This effect is most significant when environmental stochasticity is moderate. For example, when 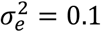, populations with 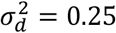 have a survival probability about 20% higher than populations with 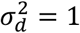, and populations with no demographic stochasticity can have even much higher survival probability. However, the difference decreases when environmental stochasticity is so strong that the population is very likely to go extinct.

## Discussion

In this paper, we demonstrate that the type of stochasticity that a population experiences can play a key role in determining both whether, and how, a population successfully undergoes evolutionary rescue. Specifically, whether stochasticity is demographic or environmental has wide-ranging repercussions on how selection intensity and genetic variance interact to determine whether a population can evolve away from extinction. When demographic stochasticity is the dominant force, we find that a slow initial decline of the genetic variance is critical for later population persistence. Tracking of the historical dynamics shows that populations which survive the rescue process maintain a relatively high level of genetic variance, while those that go to extinction tend lose their genetic variance in the first few generations. The maintenance of genetic variance leads to ultimate persistence due to a quicker evolution towards the new phenotypic optimum. In contrast, when populations are mainly subject to environmental stochasticity, genetic variance no longer plays a crucial role in whether a population survives. Instead, it is populations that happen to have a higher initial growth rate during the environmental fluctuation that are able to persist. In fact, under environmental stochasticity, there is not much difference in the temporal dynamics of genetic variance between populations that survive and those that go extinct, which may justify the assumption of a constant genetic variance in an SDE model.

Since populations often suffer both types of stochasticity in the wild, we would expect a slower decline of genetic variance and a higher initial growth rate to both contribute to successful survival in practice. Which factor is more important will depend on the relative strength of demographic and environmental stochasticity. When there is only demographic stochasticity, the strength of stochasticity has little influence on the survival probability (e.g. Fig. 6). However, with the addition of environmental stochasticity, even a slight decrease in the strength of demographic stochasticity can greatly increase the survival probability. The reason is that when there is environmental stochasticity, due to fluctuation in mean fitness, even a difference of only one individual can greatly influence the later fate of a population (see, e.g., Appendix IV). Therefore, as the population shrinks to a small size and demographic stochasticity dominates the variation, a slight difference in the strength of demographic stochasticity can make a large difference in the survival probability.

Genetic variance is often considered beneficial to the survival of populations during adaptation. However, many studies predict that the level of genetic variance should be decreased under stabilizing selection, because it can impose a fitness load in this situation (Bull 1987, Lande and Shannon 1996, Kawecki 2000). This is not a contradiction, but rather suggests that the effects of genetic variance should be discussed within a well-defined context. When populations have the same phenotypic variance, a larger genetic variance always enhances the probability of persistence, practically, the time when the observation is made is of importance (see also Martin et al. 2013). Under strong selection, populations either adapt quickly and survive, or go extinct quickly, and thus higher genetic variance has the most prominent benefit on the population survival probability early in the evolutionary rescue process, while when selection is weak it is better to compare the survival probabilities in later generations to see the effects of selection intensity on survival. On the other hand, when populations have similar environmental variance, genetic variance is a double-edged sword and does not always enhance survival. There exists a “heritability optimum” that maximizes the survival probability at a given time. The level of this heritability optimum increases quickly at first and gradually decreases and converges to a certain level over time, provided that the environmental shift is not too small. This pattern indicates that within the two antagonistic effects caused by genetic variance, a fast evolution rate plays a more important role in the early stage of the rescue process, while a lower fitness load has more importance effects on survival over the longer term.

The above results assume that genetic variance is invariant over time. While this assumption is widely assumed in quantitative genetic models (Gomulkiewicz et al. 1995, Lande and Shannon 1996, Chevin et al. 2010), in our simulation, we see a decrease of genetic variance during evolutionary rescue (e.g., Fig. 5). Therefore, populations with a higher level of genetic variance in nature may enjoy a higher survival probability for a longer time than predicted by our analytical model, which assumes a constant genetic variance. In fact, we find that adopting the assumption of a constant genetic variance overestimates the survival probability in early generations but underestimates it in later generations (Fig. S5). Since selection lowers the genetic variance, selection itself may automatically confer benefits for populations to better track the heritability optimum curve (Burger and Lynch 1995). A mechanistic model is called for to incorporate both genetic and demographic changes during evolutionary rescue.

As discussed above, the effects of genetic variance on the survival probability depend on time. Therefore, in future empirical studies, the tracking of how genetic variance affects the population survival probability over generations should provide more insights into the evolutionary rescue process. Note that although populations show different genetic and demographic dynamics under demographic and environmental stochasticity, we found the effects of genetic variance on the survival probability to be robust, simplifying the application of these results to empirical cases where the contributions of these types of stochasticity are unknown.

Apart from our genetic assumptions, our model also made several assumptions about demographic growth. The effects of density on the growth rate are not captured. Therefore, the survival probability might be overestimated, because in nature the growth of a population is expected to be slower as population size increases. This effect may be most prominent under a low carrying capacity (Boulding et al. 2001, Anciaux et al. 2019). Also, we assume that the strengths of demographic or environmental stochasticity (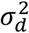 and 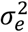) are constant in our stochastic equation (4). However, the SDE method is an approximation of the birth-death model, which is a more mechanistic way to model reality. In a birth-death model, it has been shown that the strength of demographic stochasticity 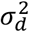 tends to change with time (Strang et al. 2019). In fact, the SDE and birth-death models are related as 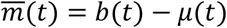 and 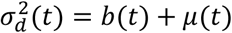, where *b*(*t*) and *μ*(*t*) are birth and death rates. Nevertheless, as the population becomes more adapted over time, we expect the birth rate *b*(*t*) to increase and the death rate *μ*(*t*) to decrease, thus 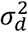 may not change dramatically, and the assumption of a constant 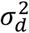 may be reasonable. However, as discussed in Appendix II, the strength of environmental stochasticity 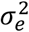 may decrease as the mean growth rate of the population increases with time, and as a result, the assumption of a constant 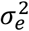 may underestimate the survival probability in the long term. If 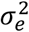 changes, this also suggests that the relative strength of demographic stochasticity versus environmental stochasticity will vary with time, which may cause the key factors that determine population persistence to be more complicated. More realistic estimates of how 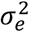 and 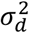 change with time during evolutionary rescue may require empirical studies.

In conclusion, this work shows that demographic and environmental stochasticity can have different effects on how the selection intensity and genetic variance influence the survival probability of a population, and presents the key factors that determine the fate of populations under the two types of stochasticity. Since the type of stochasticity has different evolutionary effects, our work suggests that explicitly distinguishing and measuring different types of stochasticity may be important for studies of evolutionary rescue.

**Table 1:**
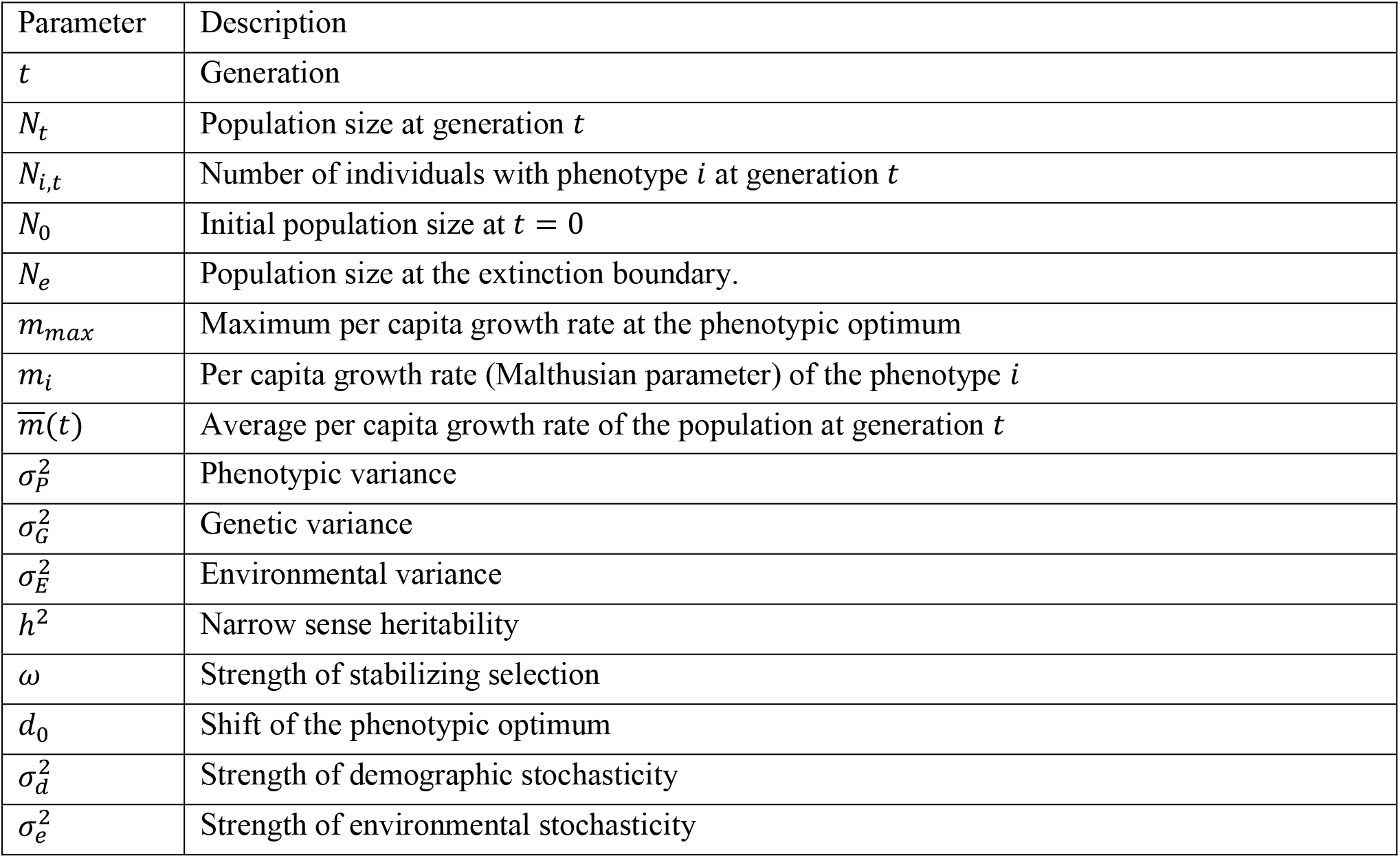
Biological meanings of symbols used in the model

| Parameter | Description |
| --- | --- |
| $t$ | Generation |
| $N_t$ | Population size at generation $t$ |
| $N_{i,t}$ | Number of individuals with phenotype $i$ at generation $t$ |
| $N_0$ | Initial population size at $t = 0$ |
| $N_e$ | Population size at the extinction boundary. |
| $m_{max}$ | Maximum per capita growth rate at the phenotypic optimum |
| $m_i$ | Per capita growth rate (Malthusian parameter) of the phenotype $i$ |
| $\overline{m}(t)$ | Average per capita growth rate of the population at generation $t$ |
| $\sigma_P^2$ | Phenotypic variance |
| $\sigma_G^2$ | Genetic variance |
| $\sigma_E^2$ | Environmental variance |
| $h^2$ | Narrow sense heritability |
| $\omega$ | Strength of stabilizing selection |
| $d_0$ | Shift of the phenotypic optimum |
| $\sigma_d^2$ | Strength of demographic stochasticity |
| $\sigma_e^2$ | Strength of environmental stochasticity |

## Acknowledgments

We would like to thank Brian Lerch for very helpful comments on an earlier draft of this paper, Joel Kingsolver for the suggestions on the model analysis, the editor and two anonymous reviewers for offering intuitive interpretations of the results.

## Supplementary Materials

## Appendix I: Definition of the per capita growth rate under Itô and Stratonovich calculi

In this section, we first briefly discuss the definition of the per capita gowth rate *m_i_* under the two types of calculi for stochastic differential equations. After that, we prove that when *m_i_* is defined as the *artihemetic mean growth rate*, the per capita growth rate of the total population 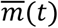 is still the arithmetic mean.

Although the changes of population size usually happen in a discrete manner, they are often approximated with stochastic differential equations. However, depending on the interpretation of the noise term, there are two major types of calculi that can be used: Itô and Stratonovich (Turelli 1977, Klebaner 2012). In this paper, we use *σdB_t_* and *σ* ° *dB_t_* to separately denote Itô and Stratonovich calculus. Actually, whether to use Stratonovich or Itô calculus depends on the biological meaning of the average per capita growth *m*(*t*) (See Turelli 1977, Braumann 2007 for detailed discussion). As proved by Braumann, for the SDE under Itô calculus:

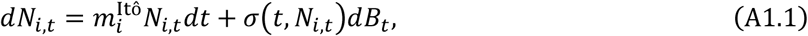

where the per capita growth rate 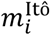 is actually the *arithmetic average growth rate*, explicitly defined as:

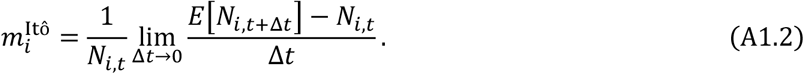

Here *E*[*N*_*i,t*+Δ*t*_] is the expected population size at time *t* + Δ*t*. For the SDE under Stratonovich calculus:

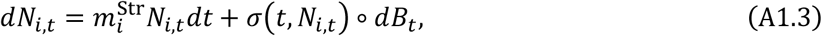

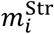 is actually the *geometric average growth rate*, defined as:

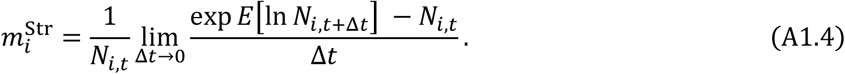

It should be noted that usually 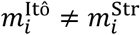. Therefore, whether to use Itô or Stratonovich calculus depends on the definition and measurement of the growth rate *m_i_*. In the following paragraph, we will show that using the *arithmetic average growth rate* is more convenient because it is additive.

Note that we want to model the growth of the total population number *N_t_*. In a deterministic model, the population dynamics of individuals with phenotype *i* is *dN_i,t_* = *m_i_N_i,t_dt*. If the dynamics of the total population size is described by 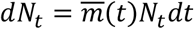, it can be easily proven that the growth rate of the total population is just the arithmetic average across all the individuals as:

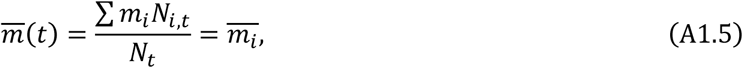

However, in stochastic equations, if *m_i_* is defined as the *arithmetic average growth rate* of *N_i,t_*, i.e., the number of indivduals with phenotype *i*, it is not naturally guaranteed that 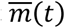 calculated in equation (A1.5) is still the *arithmetic average growth rate* of the total population size *N_t_*. Here we prove that this is the case. In fact, from equation (A1.2), it can be easily seen that the expected number of individuals with phenotype *i* at time *t* + Δ*t* is 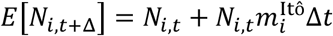, given small Δ*t*. Therefore, the expectation of the total population size at time *t* + Δ*t* is:

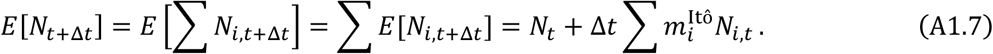

Therefore, the *arithmetic average growth rate* for *N_t_* is:

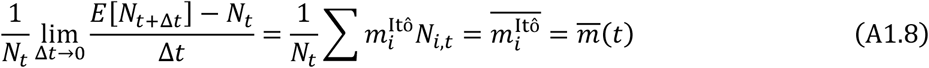

This means that we can safely write the SDE of the total population size *N_t_* as:

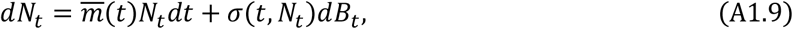

and the above equation can still be interpreted with Itô calculus. The strength of noise is just the sum of the noise of all the individuals:

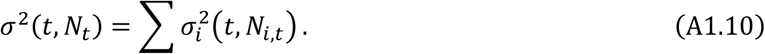

On the other hand, however, if *m_i_* is defined as the *geometric average growth rate* of *N_i,t_* (see equation (A1.4)), it is clear that the *geometric average growth rate* of the total population size *N_t_*:

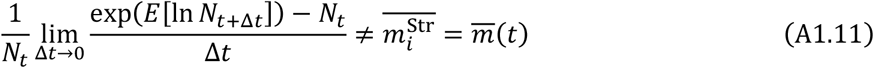

In this case, the SDE for *N_t_* cannot be written as 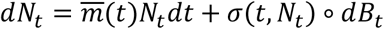. Therefore, for convenience, we define all the *m_i_* in this paper to be the *arithmetic average growth rate*, so that we can directly average the per capita growth m*i* across all the individuals and use it to describe the demographic dynamics of the total population as in equation (A1.8), where 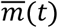 is expressed by equation (4) in the main text.

## Appendix II: The strength of environmental stochasticity 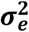 for a population under evolutionary rescue process can be large

In previous studies of wild populations, the strength of environmental stochasticity *σ_e_* is often estimated to be small in magnitude (Lande et al. 2006, Sæther et al. 1998). However, in this paper, we usually choose the value of *σ_e_* chosen to be of the order ~ 1. Here we show that the value *σ_e_* is greatly influenced by the population growth rate and explain why our choice for the value of *σ_e_* is reasonable.

As is discussed in the Methods section, the strength of demographic and environmental stochasticity 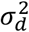 and 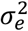 in the SDE measures the variance of population size in an infinitesimal time interval. However, in reality, the population size changes in a discrete manner, and most studies actually use data with the time interval to be months or a year. Based on equation (3), the population size at *t* + 1 is 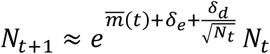, so the deviation of *N*_*t*+1_/*N_t_* from the expectation is

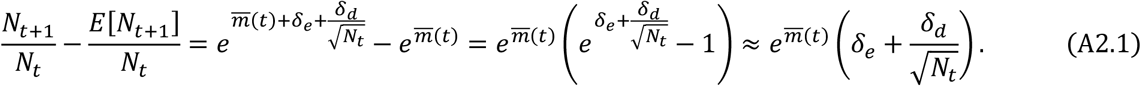

The variance of the relative population size change *N*_*t*+1_/*N_t_* is

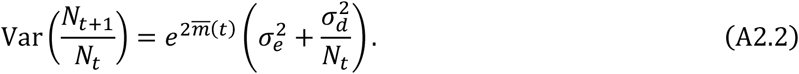

Thus 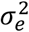 and 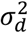 can be estimated through the value of the mean offspring number *N*_*t*+1_/*N_t_* and its variance calculated from the data, as has been done in many studies (Sæther et al. 1998, Reed et al. 2006, Reed et al. 2007). It should be noted that the estimation of 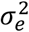 is dependent on the per capita growth rate 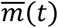. Given the same level of fluctuation in absolute fitness Var(*N*_*t*+1_/*N_t_*), when population has a negative growth rate 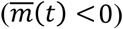, *σ_e_* can be large since 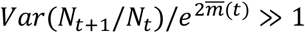. In contrast, when 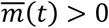, *σ_e_* should be quite small since 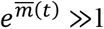. Indeed, in studies where *σ_e_* is estimated to be small, populations always have a positive growth rate. For example, in ibex populations across several habitats with the median value of *σ_e_* being about 0.1, all of the populations actually have a positive growth rate (Sæther et al. 2007). In a great tit population where 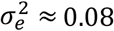, the population has a quite high growth rate of 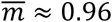 (Sæther et al. 1998). However, even in a quickly growing population, strong environmental stochasticity can occur. In another study of ibex populations, *σ_e_* is estimated to be about 0.45 even when the mean growth rate 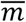 is about 0.14 (Sæther et al. 2002).

Since populations often have a negative growth rate during the early stage of the evolutionary rescue process, *σ_e_* can be large, although it should be noted that the variance of relative population size change Var(*N*_*t*+1_/*N_t_*) may also decrease as the mean fitness becomes lower., Also, we would expect 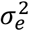 to decrease with time as 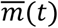 evolves to be positive, and as a result, the assumption of a constant 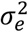 may underestimate the survival probability.

## Appendix III: Numerical simulation for population survival probability under both types of stochasticity

Under environmental stochasticity, the SDE becomes:

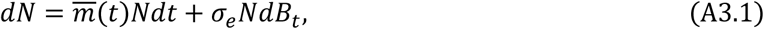

which has an explicit strong solution (Braumann 2007):

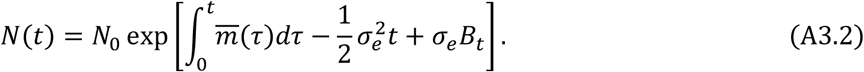

From the solution, it is easy to see the population size will never reach 0. Biologically, this is due to the fact that environmental stochasticity only influences the intrinsic growth rate *m*(*t*), rather than the population size itself. However, there is no general analytical solution for the survival probability. Therefore, we numerically calculate the survival probability using the simulation described in the following paragraph.

Here we describe the general simulation algorithm to obtain the population survival probability when both demographic and environmental stochasticity are present. The SDE for the population dynamics in this case is:

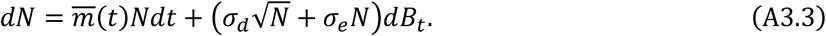

Since the above equation is interpreted under Itô calculus, we can directly discretize the continuous process into a difference stochastic equation by a time step Δ*t*:

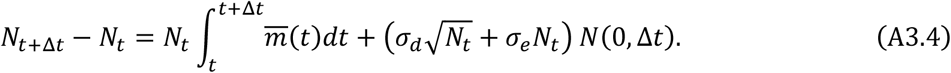

For the integral of *m*(*t*) in the above equation, we use the midpoint rule as 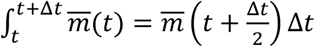. *N*(0, Δ*t*) is a normal distribution with a mean 0 and a variance Δt. For each simulation, the stochastic process starts from the initial population size *N*_0_, and stops whenever it reaches the lower boundary *N_e_*. For simulations throughout the paper, we choose Δ*t* = 0.01, and the population survival probability under a certain set of parameter values is calculated from 10^5^ replications. The program for this numeric solution is wirtten in R and is available on Dryad.

## Appendix IV: Individual-based simulation

**Table S1:**
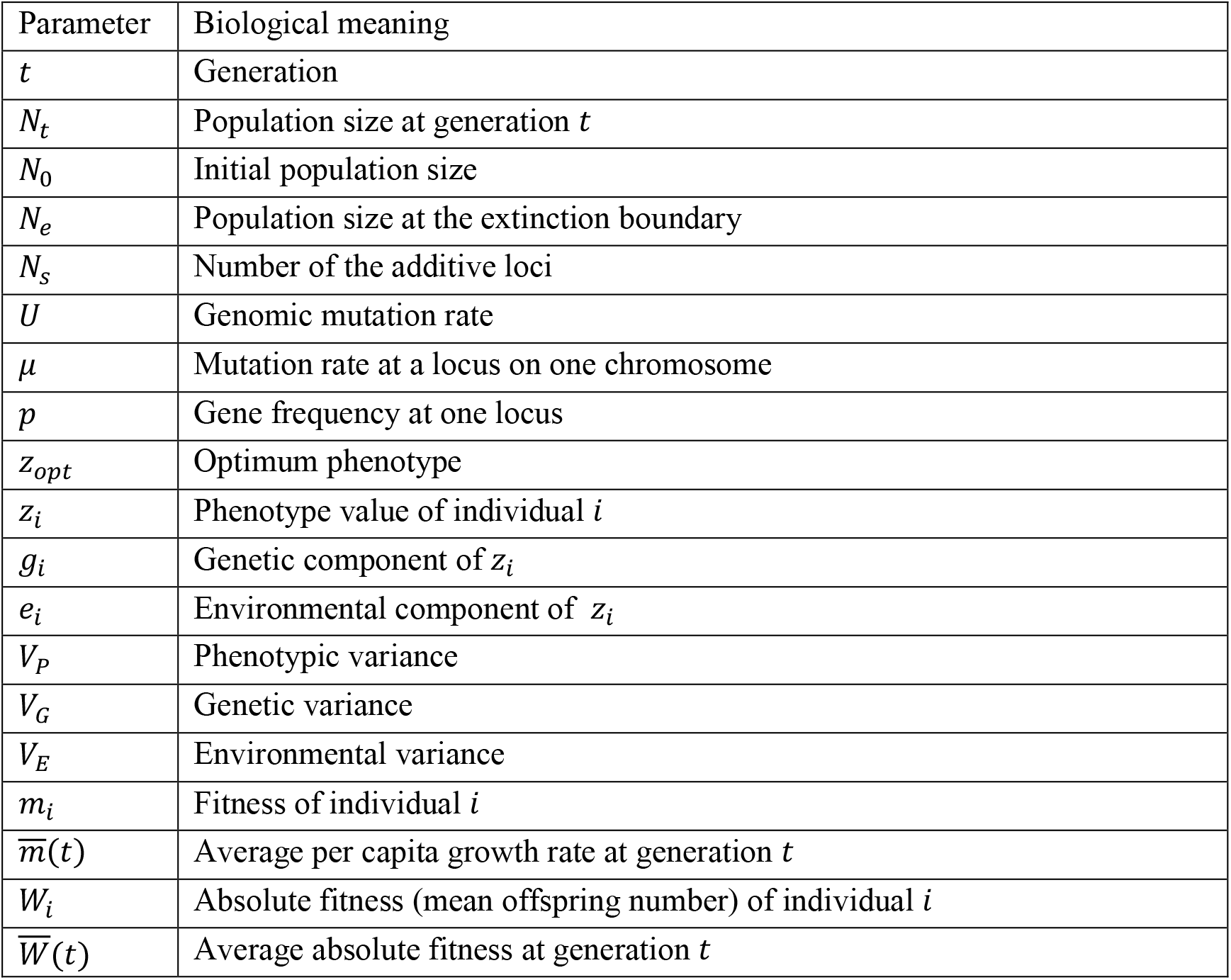
List and description of all parameters used in simulation

The simulation algorithm consists of a genetic and a demographic component. Our genetic assumptions follow the algorithm of the previous evolutionary rescue study by Boulding et al. 2001. The notation used in the simulation is listed in Table S1. We assume a diploid, hermaphroditic population. The individual phenotype being selected upon is controlled by *N_s_* addtitive loci with free recombination and identical effects. If each allele has a phenotypic effect of 1, then the possible phenotypic value has a range of [0,2*N_s_*]. Every generation, these loci have an average genomic mutation rate of *U*, which means a mutation rate of *μ* = *U*/2*N_s_* at each locus on one chromosome. The individual phenotypic value can be written as:

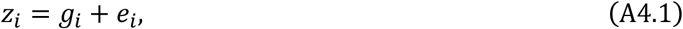

where *g_i_* is the addtive genetic contribution and *e_i_* is the environmental component, which is drawn from a zero-mean Gaussian distribution with the variance equal to the environmental variance 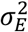. Since different values of *N_s_* result in different ranges of additive values, we need to scale the additive genetic effects. Since we assume there is no linkage between loci, when the gene frequency at each locus is *p* = 0.5 the genetic variance is at its maximum 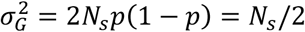. Therefore, for each simulation, we first set *V_P_* and *V_E_* to be some constant number of units, and then scale the additive effects of each allele by dividing by the scaling number 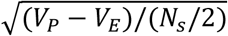. However, it should be noted that due to drift and mutation, the actual *V_G_* is always smaller than the maximum value *V_P_* – *V_E_*.

The simulation is divided into two parts: the initialization stage and evolutionary rescue stage. The initialization stage generates a population at the selection-mutation-drift equilibrium. During this stage, we set the optimum phenotype to *z_opt_* = 0 units and assume that the population size is constant. We run the initialization stage for an long enough time *T_init_* to ensure the population reaches equilibrium (usually 2000 generations). After that, we let the environmental shift occur by increasing the optimum phenotype *z_opt_* to *z_opt_* = *d*_0_ model units. During the evolutionary rescue stage, the algorithm for demographic change is modified from Anciaux et al. (2019). For each replicate, the program stops when the population reaches the extinction boundary *N_e_*, or the population is considered as rescued when the size is greater than 10^5^. Since we focus on a short time scale, the simulation also stops when the time reaches the limit *T_set_*. To obtain the survival probability, after the initialization stage, we run 500 replications for the rescue stage, so that each replicate populations starts from the same genetic and demographic condition. The extinction threshold is usually set to be *N_e_* = 0 for demographic stochasticity and *N_e_* = 1 for environmental stochasticity.

A condensed description of the algorithms follows:

### Initialization stage

Set *Z_opt_* = 0. **While (***t* > *T_init_*)

1. Mutation process: Draw the number of mutations for each individual from a binomial distribution with rate *μ*.
2. Individual phenotype and fitness: Generate the phenotype of each individual *z_i_* = *g_i_* + *e_i_*. An individual’s growth rate *m_i_* is caculated based on the fitness landscape function (equation (1) in the main paper).
3. Draw offspring genotypes: For each offspring, the parent is randomly drawn from the previous population with a probability proportional to the relative fitness *W_i_/W_max_*, where *W_i_* = *e^m_i_^* is the absolute fitness of an individual with phenotype *i*, and *W_max_* is the maximum absolute fitness in the parental population. **end while**

### Evolutionary rescue stage for demographic stochasticity

Increase *z_opt_* to *d*_0_. **While (**COND = **if** ((*N_t_* > 10^5^) or (*t* > *T_set_*)))

1. Mutation process: Draw the number of mutations for each individual from a binomial distribution with rate *μ*.
2. Individual phenotype and fitness: Generate the phenotype of each individual *z_i_* = *g_i_* + *e_i_*, and calculate each individual’s growth rate 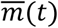. After that, calculate the average per capita growth rate m(t) and the mean absolute fitness 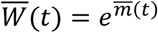.
3. Offspring number: The expected offspring number in the next generation is 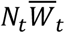. However, for demographic stochasticity, *N*_*t*+1_ is drawn from a Poisson distribution 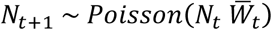, which ensures the strength of demographic stochasticity to be approximately a constant 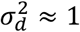.
4. Offsping genotypes: For each offspring, the two parents are randomly drawn from the previous population with a probability proportional to the relative fitness *W_i_/W_max>_*. For each parent, the gamete is randomly drawn after recombination.
5. **if** (*N_t_* ≤ *N_e_*): **break**; **end while**

### Evolutionary rescue stage for environmental stochasticity

Increase *z_opt_* to *d*_0_. **While (**COND = **if (** (*N_t_* > 10^5^) or (*t* > *T_set_*)))

1. Mutation process: Draw the number of mutations for each individual from a binomial distribution with rate *μ*.
2. Individual phenotype and fitness: Generate the phenotype of each individual, *z_i_* = *g_i_* + *e_i_*, and calculate *m_i_* and 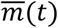. To model environmental stochasticity, a random number *δ* is drawn from the normal distribution 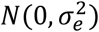. Therefore, the average per capita growth rate becomes 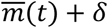, and the corresponding mean absolute fitness is 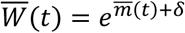.
3. Offspring number: Since the expected offspring number 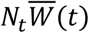 is numeric, it needs to be integrated. Two functions are chosen: 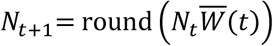 and 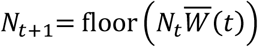.
4. Offspring genotypes: For each offspring, the two parents are randomly drawn from the previous population with a probability proportional to the relative fitness *W_i_/W_max_*. For each parent, the gamete is randomly drawn after recombination.
5. **if** (*N_t_* ≤ *N_e_*): **break**; **end while**

For environmental stochasticity, it is not easy to make a sensible comparison with the SDE model predictions, due to the intrinsic limitations of the simulation algorithm. In fact, as shown in Fig. S1, the two integrating functions round and floor can yield very different results. The difference in survival rate between the functions becomes smaller as environmental stochasticity 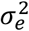 increases (*σ_e_* = 1 in Fig. S1(c)).

To explain this phenomenon, we further examined the genetic and demographic dynamics. We find no difference between the historical dynamics of the genetic variance and mean growth rate across the replicate populations under the two functions. However, it is when the population size declines to a low level that the two functions start to diverge markedly. Figs. S2(a) and S2(b) respectively show the demographic dynamics of those populations that successfully survive through the time period of 100 generations under the round and floor functions. In both figures, the population size quickly declines to only few individuals after the shift happens. However, under the round function, populations are allowed to reach a lower boundary than under a floor function. When the expected population size in the next generation 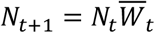 is a numeric value, the round function causes *N*_*t*+1_ to have about a 50% probability of going one step up and a 50% probability of going one step down, while a floor function always causes *N*_*t*+1_ to go one step down. When the population size is large, this difference of about 0.5 individual is not important. Nevertheless, it really matters when the population size is close to the edge of the extinction boundary *N_e_ = 1*, as even a difference of only one individual can greatly help a population to escape from the absorbing boundary.

## Supplementary Figures

**Figure S1.**
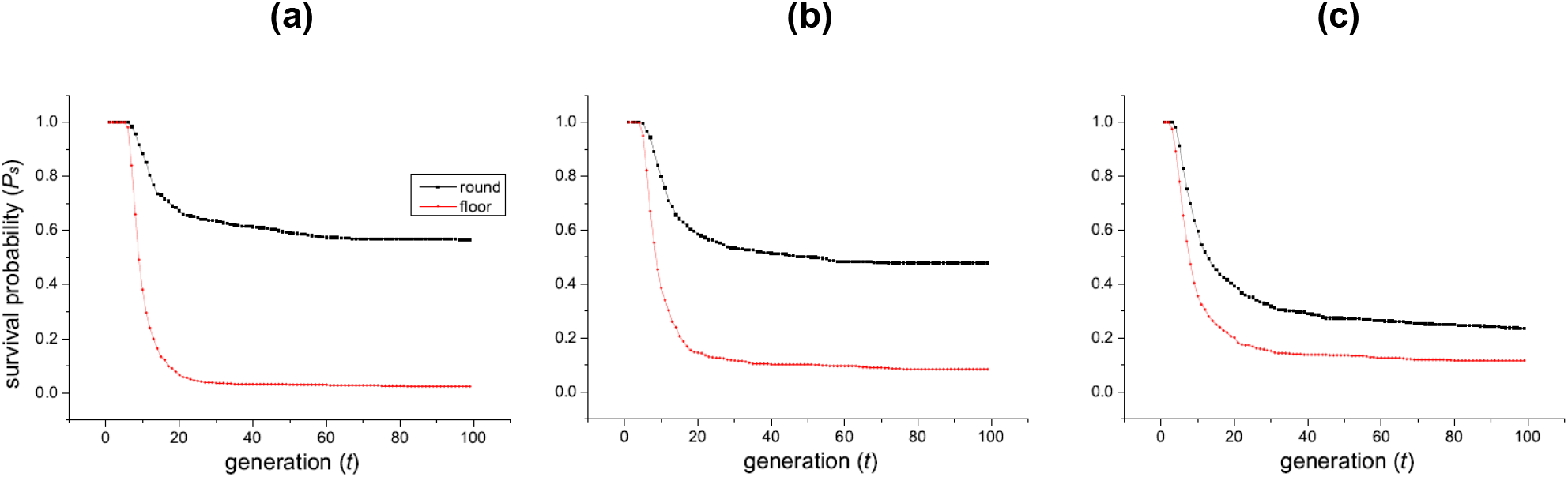
Comparison of the population survival probability dynamics between the two functions round and floor during the environmental stochasticity simulation. The strength of environmental stochasticity is 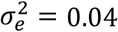 for panel (a), 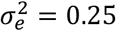 for (b), and 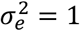 for (c). The black line “round” shows the results given by the round function, and the red line “floor” is from the floor function. Under weak environmental stochasticity (panel (a)), the round function gives much higher survival rate than floor. As *σ_e_* gets larger, the two lines becomes closer to each other. Parameter values used during the simulation are *N*_0_ = 500, *N_e_* = 1, *m_max_* = 0.2, *ω* = 0.3, *d*_0_ = 3, *V_P_* = 1, *V_E_* = 0.5, *U* = 0.5.

**Figure S2.**
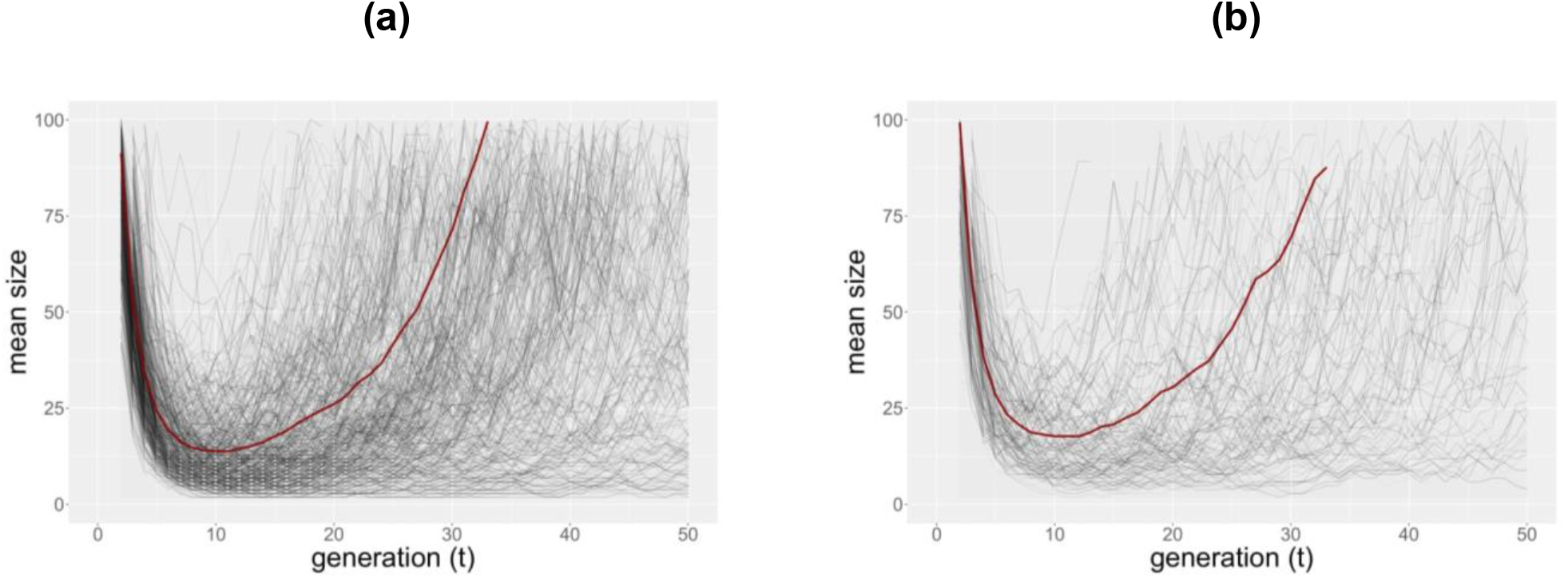
Demographic trajectories across replicate populations that survive through the period of 100 generations under the round (panel (a)) and floor (panel (b)) function. Demographic dynamics of the replicate populations are colored from grey to black, while the red line shows the average population size. There are more lines in panel (a) because the round function gives a higher survival rate. Generally, a round function allows the populations to reach a lower boundary without going extinct. This is also reflected in the red curve of the average population size, as populations under a floor function generally have a slightly higher average size. which can be seen by comparing the minimum of the red curves in panels (a) and (b). Environmental stochasticity is set to be 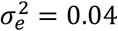. Parameter values are *N*_0_ = 500, *N_e_* = 1, *m_max_* = 0.2, *ω* = 0.3, *d*_0_ = 3, *V_P_* = 1, *V_E_* = 0.5, *U* = 0.5.

**Figure S3.**
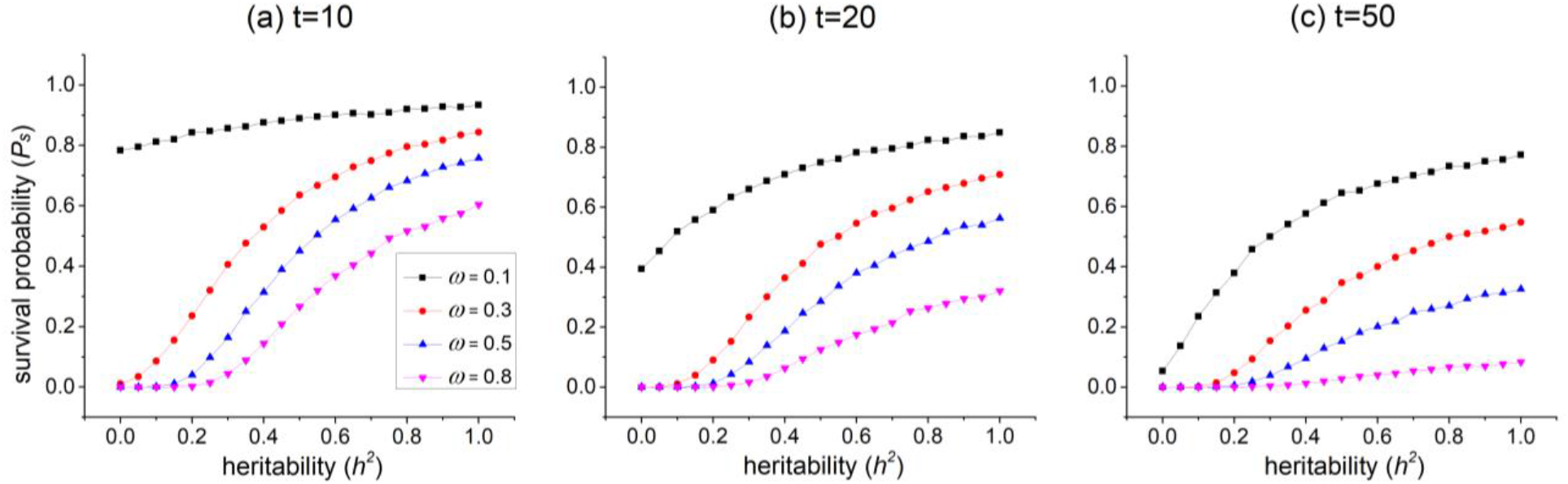
Changes of the survival probability with heritability *h*^2^ under environmental stochasticity given a fixed phenotypic variance. Panels (a), (b), and (c) respectively show snapshots at the three time points *t* = 10,20, and 50. In each panel, the colored lines indicate different selection intensities at *ω* = 0.1,0.3,0.5, and 0.8. Generally, under stronger selection, the most prominent change of over *h*^2^ happens earlier in time. For *ω* = 0.8, the most prominent change appears at time *t* = 10, while for *ω* =0.1, heritability has the most significant effect on the population survival rate at *t* = 50. However, unlike demographic stochasticity in Fig. 2, under weak selection (*ω* = 0.1), there is still a risk of extinction even under high level of heritability at time *t* = 50. Parameters for all three panels are 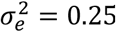, *N*_0_ = 500, *d*_0_ = 3, *m*_0_ = 0.2, 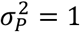.

**Figure S4.**
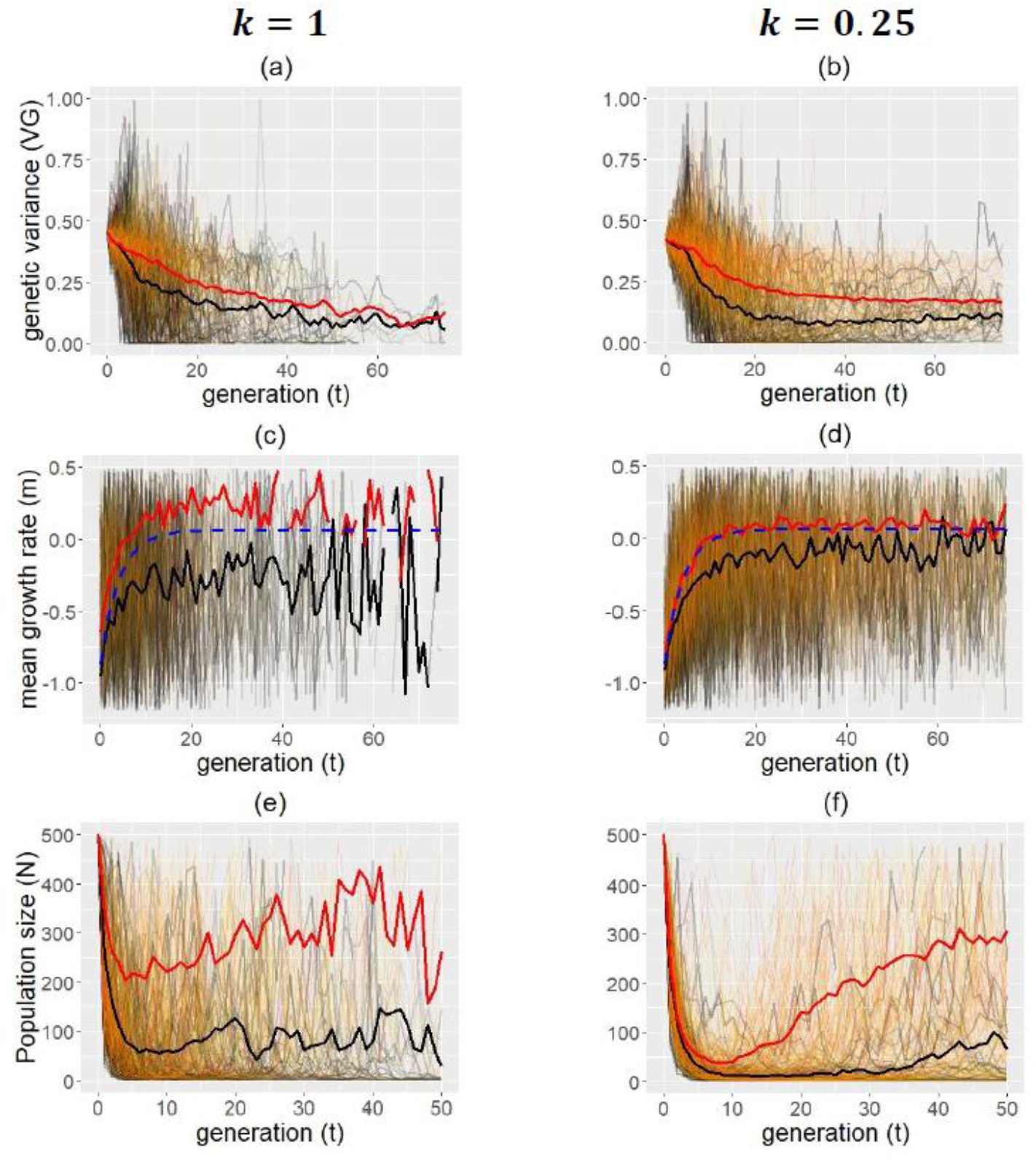
Genetic and demographic dynamics when the strength of environmental stochasticity is proportional to the absolute fitness (i.e., 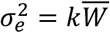). Other parameters used are the same as those in Fig. 5.

**Figure S5.**
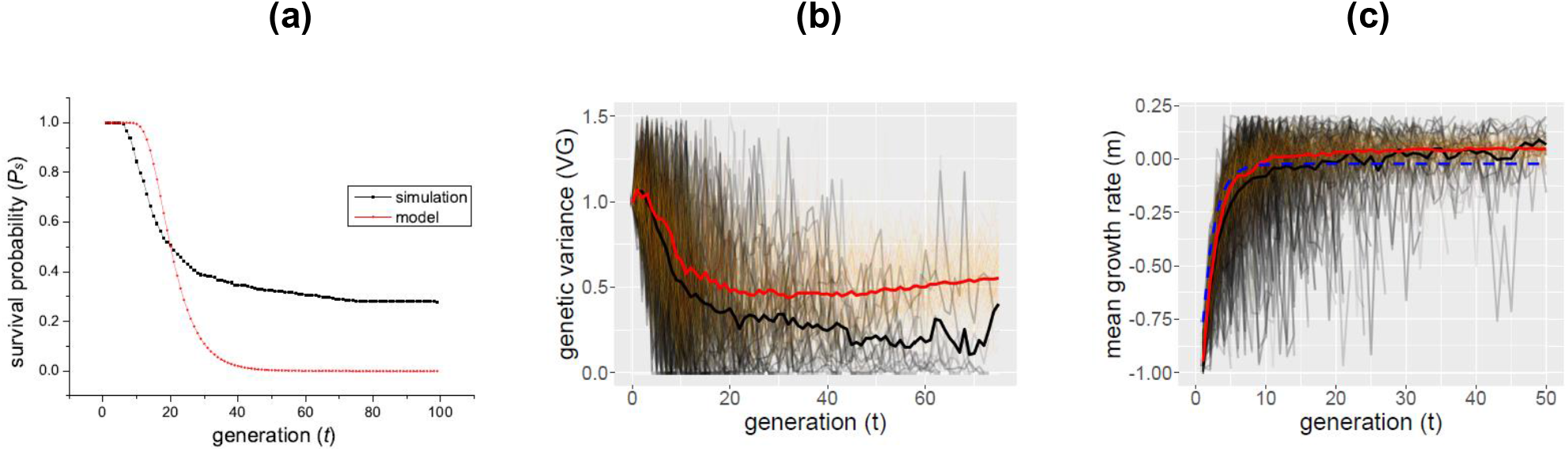
Comparison between model predictions and simulation under demographic stochasticity with strength 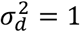. Panel (a) shows the survival probability given by the model and simulation. The model gives an overestimation before generation *t* =20, while underestimates the survival rate in the later generations. Panel (b) and (c) respectively show the dynamics of genetic variance and mean growth rate in the simulation across 500 replicate populations, with the average across the survival and extinction groups illustrated by the red and black line. The dashed line in panel (c) shows the fitness dynamics under the assumption of a constant genetic variance. It can be seen that the model predicts a quicker evolution and a higher fitness than the simulation before about *t* = 10. However, both the survival and extinction groups later have a higher fitness than the model prediction due to the decline in their genetic variance. Parameter values used in the simulation are that *N*_0_ = 500, *N_e_* = 0, *m_max_* = 0.2, *ω* = 0.3, *d*_0_ = 3, *V_P_* = 2, *V_E_* = 0.5, *U* = 0.5, and the value of genetic variance used in the model is 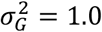, which is the initial value of the populations in simulation.

